# Joint structural annotation of small molecules using liquid chromatography retention order and tandem mass spectrometry data

**DOI:** 10.1101/2022.02.11.480137

**Authors:** Eric Bach, Emma L. Schymanski, Juho Rousu

## Abstract

We present LC-MS^2^Struct, a machine learning framework for structural annotation of small molecule data arising from liquid chromatography-tandem mass spectrometry (LC-MS^2^) measurements. LC-MS^2^Struct jointly predicts the annotations for a set of mass spectrometry features in a sample, using a novel structured prediction model trained to optimally combine the output of state-of-the-art MS^2^ scorers and observed retention orders. We evaluate our method on a dataset covering all publicly available reversed phase LC-MS^2^ data in the MassBank reference database, including 4327 molecules measured using 18 different LC conditions from 16 contributors, greatly expanding the chemical analytical space covered in previous multi-MS^2^ scorer evaluations. LC-MS^2^Struct obtains significantly higher annotation accuracy than earlier methods and improves the annotation accuracy of state-of-the-art MS^2^ scorers by up to 106%. The use of stereochemistry-aware molecular fingerprints improves prediction performance, which highlights limitations in existing approaches and has strong implications for future computational LC-MS^2^ developments.

## Introduction

Structural annotation of small molecules in biological samples is a key bottleneck in various research fields including biomedicine, biotechnology, drug discovery and environmental sciences. Samples in untargeted metabolomics studies typically contain thousands of different molecules, the vast majority of which remain unidentified [1–3]. Liquid chromatography (LC) tandem mass spectrometry (LC-MS^2^) is one of the most widely used analysis platforms [4], as it allows for high-throughput screening, is highly sensitive and applicable to a wide range of molecules. In LC-MS^2^, molecules are first separated by their different physicochemical interactions between the mobile and stationary phase of the LC, resulting in retention time (RT) differences. Subsequently, they are separated according to their mass-to-charge ratio in a mass analyzer (MS^1^). Finally, the molecular ions are isolated and fragmented in the tandem mass spectrometer (MS^2^)

For each ion, the recorded fragments and their intensities constitute the MS^2^ spectrum, containing information about the substructures in the molecule and serves as a basis for annotation efforts. In typical untargeted LC-MS^2^ workflows, thousands of MS features (MS^1^, MS^2^, RT) arise from a single sample. The goal of structural annotation is to associate each feature with a candidate molecular structure, for further downstream interpretation.

In recent years, many powerful methods [5, 6] to predict structural annotations for MS^2^ spectra have been developed [7, 11–18, 8–10]. In general, these methods find candidate molecular structures potentially associated with the MS feature, for example, by querying molecules with a certain mass from a structure database such as HMDB [19] or PubChem [20] and subsequently computing a match score between each candidate and the MS^2^ spectrum. The highest scoring candidate is typically considered as the structure annotation of a given MS^2^. Currently, even the best-of-class methods only reach an annotation accuracy of around 40% [9] in evaluations when searching large candidate sets such as those retrieved from PubChem. Therefore, in practice, a *ranked list* of molecular structures is provided to the user (*e*.*g*. top 20 structures). This level of performance is still a significant hindrance in metabolomics and other fields.

Interestingly, RT information remains underutilized in automated approaches for structure annotation based on MS^2^, despite RTs being readily available in all LC-MS^2^ pipelines and generally recognized as contributing valuable information [21, 22]. An explanation is that a molecule generally has different RTs under different LC conditions (mobile phase, column composition, etc.) [23, 24]. Typically, the RT information is used for post-processing of candidate lists, *e*.*g*. by comparing measured and reference standard RTs [3, 24]. This approach, however, is limited by the availability of experimentally determined RTs of reference standards. RT prediction models [25, 24], on the other hand, allow to predict RTs based solely on the molecular structure of the candidate, and have been successfully applied to aid structure annotation [14, 26–29]. However, such prediction models generally have to be calibrated to the specific LC configuration [3], requiring at least some amount of target LC reference standard RT data to be available [21, 30, 29]. Recently, the idea of predicting retention *orders* (RO), *i*.*e*., the order in which two molecules elute from the LC column, has been explored [31–34]. ROs are largely preserved within a family of LC systems (*e*.*g*. reversed phase or HILIC systems). Therefore, RO predictors can be trained using a diverse set of RT reference data, and applied to out-of-dataset LC setups [31]. Integration of MS^2^ and RO based scores using probabilistic graphical models improved the annotation performance in LC-MS^2^ experiments [34].

Another somewhat neglected aspect in automated annotation pipelines is the treatment of stereochemistry, *i*.*e*. the different 3D variants of the molecules. The general assumption has been that LC-MS^2^ data does not contain sufficient information to separate stereoisomers in samples [5, 24]. As a result, MS^2^ scorers typically disregard the stereochemical information in the candidate structures and often output the same matching for different stereoisomers (*c*.*f*. [7, 9]). However, stereoisomers that vary in their double-bond orientation (*e*.*g. cis-trans* or *E-Z* isomerism) may have different shapes and thus exhibit different fragmentation and/or interactions with the LC system. Thus, ignoring stereochemistry in candidate processing may disregard LC-relevant stereochemical information. Furthermore, it is known that certain stereochemical configurations occur more frequently than others in nature and hence in the reference databases. Making use of such information can potentially improve annotation performance.

In this study we set out to provide a new perspective on jointly using MS^2^ and RO combined with stereochemistry-aware molecular features for the structure annotation of LC-MS^2^ data. We present a novel machine learning framework called LC-MS^2^Struct, which learns to optimally combine the MS^2^ and RO information for the accurate annotation of a sequence of MS features. LC-MS^2^Struct relies on the Structured Support Vector Machine (SSVM) [35] and Max-margin Markov Network [36] frameworks. In contrast to the previous work by Bach et al. [34], our framework does not require a separately learned RO prediction model. Instead, it optimizes the SSVM parameters such that the score margin between correct and any other sequence of annotations is maximized. This way, LC-MS^2^Struct learns to optimally use the RO information from a set of LC-MS^2^ experiment. We trained and evaluated LC-MS^2^Struct on all available reversed phase LC data from MassBank (MB) [37], including a combined total of 4327 molecules from 18 different LC configurations, hence reaching unprecedented measurement diversity in the model evaluation. Our framework is compared with three other approaches: RT filtering, log*P* predictions [14], and RO predictions [34]. LC-MS^2^Struct can be combined with any MS^2^ scorer, and is demonstrated with CFM-ID [12, 10], MetFrag [14, 7] and SIRIUS [11, 9]. The use of chirality encoding circular molecular fingerprints [38] in the predictive model allows to distinguish and rank different stereoisomers based on the observed ROs.

## Overview of LC-MS^2^Struct

### Input and output

We consider a typical data setting in an untargeted LC-MS^2^ based experiments, after pre-processing such as chromatographic peak picking and alignment (Figure 1**a**). Such data comprises a sequence of MS features, here indexed by *σ*. Each feature consists of MS^1^ information (e.g. mass, adduct and isotope pattern), LC retention time (RT) *t*_*σ*_ and an MS^2^ spectrum *x*_*σ*_. We assume that a set of candidate molecules 𝒞_*σ*_ is associated with each MS feature *σ*. Such a set can be, for example, generated from a structure database (e.g. PubChem [20], ChemSpider [39] or PubChemLite [40]) based on the ion’s mass, a suspect list, or an in silico molecule generator (e.g. SmiLib v2.0 [41, 42]). We furthermore require that for MS^2^ spectrum *x*_*σ*_, a matching score *θ*(*x*_*σ*_, *m*) with its candidates *m* ∈ 𝒞_*σ*_ is pre-computed using an in silico tool, such as CFM-ID [12, 10], MetFrag [14] or SIRIUS [11, 9]. LC-MS^2^Struct predicts a score for MS feature *σ* and each associated candidate *m* ∈ 𝒞_*σ*_ based sequence of spectra 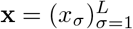, of length *L*, and the ROs derived from the observed RTs 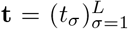. These scores are used to rank the molecular candidates associated with the MS features (Figure 1**b**).

**Figure 1:**
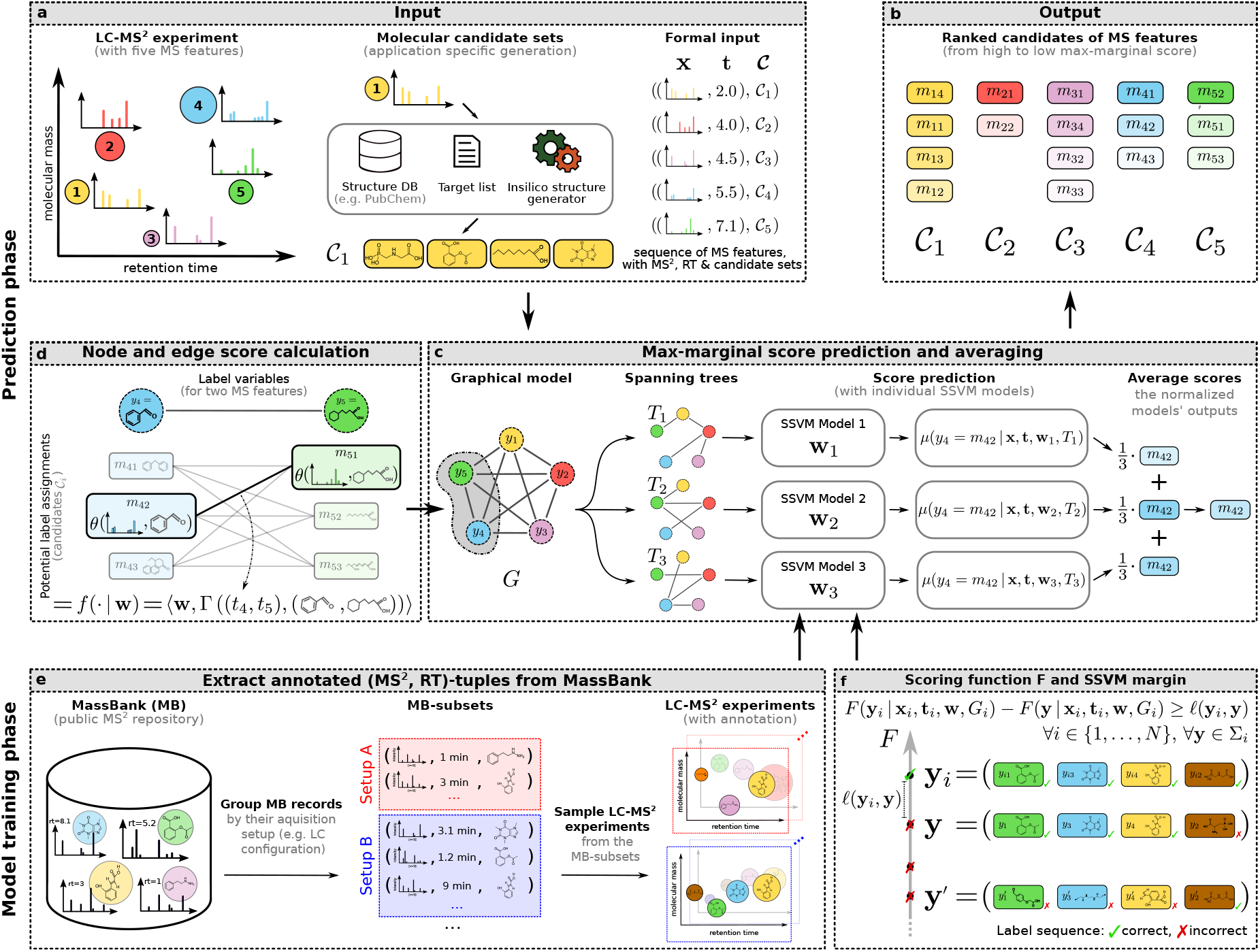
Overview of the LC-MS^2^Struct workflow. **a**: Input to LC-MS^2^Struct during the application phase. The LC-MS^2^ experiment results in a set of (MS^2^, RT)-tuples. The MS information is used to generate a molecular candidate set for each MS feature. **b**: Output of LC-MS^2^Struct are the ranked molecular candidates for each MS feature. **c**: A fully connected graph *G* models the pairwise dependency between the MS features. Using a set of random spanning trees *T*_*k*_ and Structured Support Vector Machines (SSVM) we predict the max-marginal scores for each candidate used for the ranking. **d**: The MS^2^ and RO information is used to score the nodes and edges in the graph *G*. **e**: To train the SSVM models and evaluate LC-MS^2^Struct, we extract MS^2^ spectra and RTs from MassBank. We group the MassBank records such that their experimental setups are matching, simulating LC-MS^2^ experiments. **f**: Main objective optimized during the SSVM training, where **y**_*i*_ ∈ Σ_*i*_ is the ground truth label sequence of example *i* and **y, y**^′^ ∈ Σ_*i*_ are further possible label sequences.

### Candidate ranking using max-marginals

We define a fully connected graph *G* = (*V, E*) capturing the MS features and modelling their dependencies (Figure 1**c**). Each node *σ* ∈ *V* corresponds to a MS feature, and is associated with the pre-computed MS^2^ matching scores *θ*(*x*_*σ*_, *m*) between the MS^2^ spectrum *x*_*σ*_ and all molecular candidates *m* ∈ 𝒞_*σ*_. The graph *G* contains an edge (*σ, τ*) ∈ *E* for each MS feature *pair*. A scoring function *F* is defined predicting a compatibility score between a sequence of molecular structure assignments 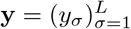 in the label-space Σ = 𝒞_1_ × … × 𝒞_*L*_ and the observed data:

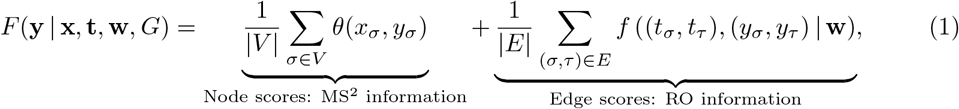

where the function *f* outputs an edge score (Figure 1**d**) expressing the agreement between the observed and the predicted RO, for each candidate assignment pair (*y*_*σ*_, *y*_*τ*_) given the observed RTs (*t*_*σ*_, *t*_*τ*_). Function *f* is parameterized by the vector **w**, which is trained specifically for each MS^2^ scorer (see next section). Using the compatibility score function *F* (Equation (1)) we compute the max-marginal scores [43] for each candidate and MS feature, defined for a candidate *m* ∈ 𝒞_*σ*_ and MS feature *σ* as the maximum compatibility score that a candidate assignment 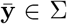 with 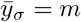 can reach:

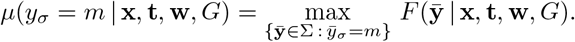

We use *μ* to rank the molecular candidates [34]. However, for general graphs *G*, the max-marginal inference problem (MMAP) is intractable. Therefore, we approximate the MMAP problem by performing the inference on tree-like graphs *T*_*k*_ randomly sampled from *G* (Figure 1**c**), for which exact inference is feasible [43, 44]. Subsequently, we average the max-marginal scores *μ*(*y*_*σ*_ = *m* | **x**_*i*_, **t**_*i*_, **w**_*k*_, *T*_*k*_) over a set of trees **T**, an approach that performed well for practical applications [45, 46, 34]. For each spanning tree *T*_*k*_, we apply a separately trained SSVM model **w**_*k*_ to increase the diversity of the predictions.

### Joint annotation using Structured Support Vector Machines (SSVM)

We propose to tackle the joint assignment of candidate labels **y** ∈ Σ to the sequence of MS features of a LC-MS^2^ experiment through structured prediction, a family of machine learning methods generally used to annotate sequences or networks [35, 47, 46]. In our model, the structure is given by the observed RO of the MS feature pairs (*y*_*σ*_, *y*_*τ*_), which provides additional information on the correct candidate labels *y*_*σ*_ and *y*_*τ*_. Given a set of annotated LC-MS^2^ experiments extracted from MassBank [37] (Figure 1**e**), we train a Structured Support Vector Machine (SSVM) [35] model **w** predicting the edge scores. SSVMs models can be optimized using the max-margin principle [35]. In a nutshell, given a set of ground truth annotated MS feature sequences, the model parameters **w** are optimized such that the correct label sequence **y**_*i*_ ∈ Σ_*i*_, that is the structure annotations for all MS features in an LC-MS^2^ experiment, scores higher than any other possible label sequence assignment **y** ∈ Σ_*i*_ (Figure 1**f**).

## Results

### Extracting training data from MassBank

Ground truth annotated MS^2^ spectra and RTs were extracted from MassBank [37], a public online database for MS^2^ data. Each individual MassBank record typically provides a rich set of meta information (see Supplementary Table 1), such as the chromatographic and MS conditions as well as molecular structure annotations. For training the SSVM model of LC-MS^2^Struct, the MassBank data was processed such that the experimental conditions were consistent *within* each MS feature set, *i*.*e*. with identical LC setup and MS configuration as in a typical LC-MS^2^ experiment, to ensure comparable RTs, ROs and MS^2^ data. We developed a Python package “massbank2db” that can process MassBank records and groups them into consistent MS feature sets, which we denote as MB-subsets. For our experiments, we sampled sequences of MS features from the MB-subsets in order to simulate real LC-MS^2^ experiments where the signals of multiple unknown compounds are measured under consistent experimental setups. Figure 1**e** illustrates the grouping and LC-MS^2^ sampling process. Two collections of MassBank data were considered: ALLDATA and the ONLYSTEREO subset.

### Comparison of LC-MS^2^Struct with other approaches

In the first experiment we compare LC-MS^2^Struct with previous approaches for candidate ranking either using only MS^2^ or additionally RT or RO information: *Only-MS*^*2*^ uses the MS^2^ spectrum information to rank the molecular candidates and serves as baseline; *MS*^*2*^*+RO* [34] uses a Ranking Support Vector Machine (RankSVM) [48, 49] to predict the ROs of candidate pairs and a probabilistic inference model to combine the ROs with MS^2^ scores; *MS*^*2*^*+RT* uses predicted RTs to remove false positive molecule structures from the candidate set, ordered by their MS^2^ score, by comparing the predicted and observed RT; *MS*^*2*^*+logP* is an approach introduced by Ruttkies et al. [14], which uses the observed RT to predict the XLogP3 value [50] of the unknown compound and compares it with the candidates’ XLogP3 values extracted from PubChem to refine the initial ranking based on the MS^2^ scores. The RO based methods (LC-MS^2^Struct and MS^2^+RO) were trained using the RTs from all available MB-subsets, ensuring that no test molecular structure (based on InChIKey first block) was used for the model training (structure disjoint). For the RT based approaches (MS^2^+RT and MS^2^+log*P*), the respective predictors were trained in a structure disjoint fashion using only the RT data available for that MB-subset. For the experiment, all MB-subsets with more than 75 (MS^2^, RT)-tuples from the ALLDATA data setup were used (see Supplementary Table 2), as the RT based approaches require LC system-specific RT training data. The ranking performance was computed for each LC-MS^2^ experiment within a particular MB-subset. The candidate molecules are identified by their InChIKey first block (*i*.*e*. the structural skeleton), hence no stereoisomers are in the candidate sets.

Each candidate ranking approach was evaluated with three MS^2^ scorers: CFM-ID 4.0 [10], MetFrag [14] and SIRIUS [9]. For LC-MS^2^Struct we use stereochemistry aware molecular fingerprints (3D) to represent the candidates.

Figure 2**a** shows the average ranking performance (top-*k* accuracy) across 350 LC-MS^2^ experiments, each encompassing about 50 (MS^2^, RT)-tuples (see Methods). LC-MS^2^Struct is the best performing method combined with any of the three MS^2^ scorers. For CFM-ID and MetFrag, LC-MS^2^Struct provides 4.7 and 7.3 percentage unit increases over the Only-MS^2^ for the top-1 accuracy, corresponding to 80.8% and 106% performance gain. In our setting, that translates to 2.4 and 3.7 additional annotations at the top rank, respectively (out of approx. 50). The performance improvement increases for larger *k*, reaching as far as 9.3 and 11.3 percentage units at top-20, which means 4.7 and 5.7 additional correct structures, respectively, in the top-20. For SIRIUS, the improvements are more modest, on average around 2 percentage units for top-1 to top-20. That might be explained by the higher baseline performance of SIRIUS. Nevertheless, SIRIUS can be improved for particular MB-subsets (see Figure 2**b** and the discussion in the next section).

**Figure 2:**
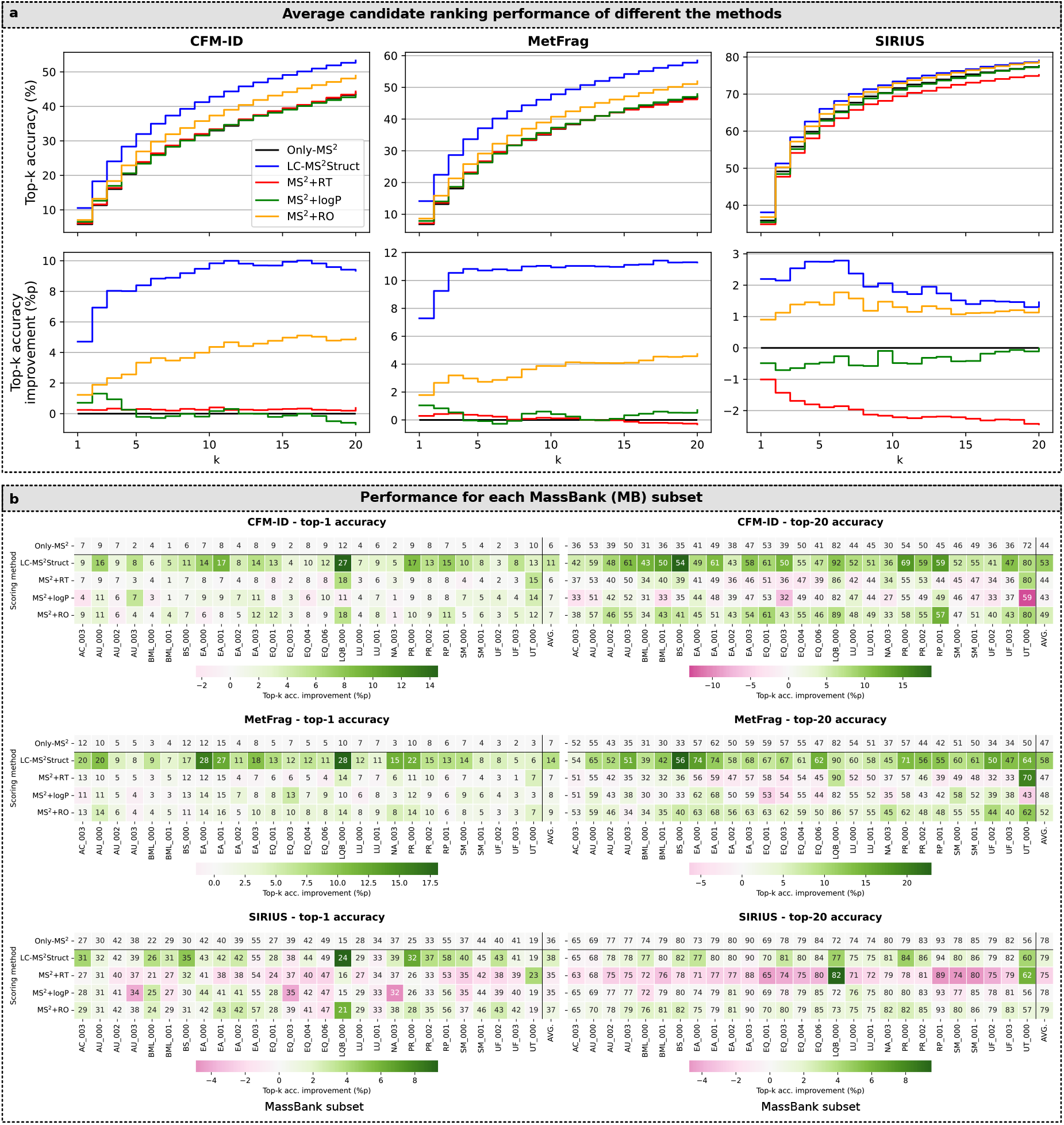
Different approaches to combine MS^2^ and retention time (RT) information: **a**: Comparison of the performance, measured by top-*k* accuracy, for the different ranking approaches combining MS^2^ and RT information. The results shown are averaged accuracies over 350 sample MS feature sequences (LC-MS^2^ experiments). **b**: Average top-*k* accuracies per MassBank (MB) subset, rounded to full integers. The color encodes the performance improvement of each score integration method compared with Only-MS^2^.

The runner-up score integration method is MS^2^+RO, which also makes use of predicted ROs. For CFM-ID and MetFrag it leads to about one-third to a half of the performance gain as LC-MS^2^Struct. The approaches relying on RTs, either by candidate filtering (MS^2^+RT) or through log*P* prediction (MS^2^+log*P*), lead to only minor improvements for MetFrag and CFM-ID, but none for SIRIUS, for which MS^2^+RT even leads to a decrease in ranking performance by about 2 percentage units. An explanation for this is that the filtering approach removes on average 4.7% of the correct candidates, which leads to false negative predictions.

The performance gain by using either RO or RT varies between the MB-subsets with differing LC-MS^2^ setups (see Supplementary Table 3) and compound class compositions (see Extended Data Figure 1). We illustrate these differences in Figure 2**b**. Applying LC-MS^2^Struct improves the ranking performance in almost all MB-subsets, including the SIRIUS MS^2^-scorer (some very slight decreases were observed in some SIRIUS scored sets). This is in stark contrast to the RT based approaches (MS^2^+RT and MS^2^+log*P*), which often lead to less accurate rankings, especially for SIRIUS. Furthermore, as seen already in the average results (Figure 2**a**), the benefit of LC-MS^2^Struct depends on the MS^2^ base scorer. For example, the top-1 accuracy of the subsets “AC 003” and “NA 003” can be greatly improved for MetFrag but show little improvement for CFM-ID. Both datasets are natural product toxins, which are perhaps poorly explained by the bond-disconnection approach of MetFrag. In contrast, for “RP_001” and “UF_003” the largest improvements (top-1) can be reached for CFM-ID. The RT filtering approach (MS^2^+RT) performs particularly well for “LQB_000” and “UT_000”. These subsets mostly contain lipids and lipid-like molecules (see Extended Data Figure 1).

Since the RT prediction models are trained using only data from the respective MB-subsets, more accurate models may be reached for less heterogeneous subsets of molecules. Hence, the RT filtering could work well in such cases [26].

### Performance analysis of LC-MS^2^Struct for different compound classifications

Next we investigate how LC-MS^2^Struct can improve the annotation across different categories in two molecule classification systems, ClassyFire [51] and PubChemLite [40]. Figure 3 shows the average top-1 and top-20 accuracy improvement of LC-MS^2^Struct over the Only-MS^2^ baseline for each ClassyFire super-class and PubChemLite annotation category. For ClassyFire (Figure 3**a**), the ranking performance improvement for the different super-classes depends on the MS^2^ scorer. For example, the top-1 accuracy of “Alkaloids and derivatives” can be improved by 10.8 percentage units for MetFrag, but improves much less for CFM-ID and SIRIUS (1.9 and 3.5 percentage units, respectively). For “Organic oxygen compounds”, in contrast, the top-1 accuracy improves by about 10 percentage units when using both CFM-ID and MetFrag, whereas only half that improvement is observed for SIRIUS. This suggests that the CFM-ID results may be improved with the inclusion of more “Alkaloids and derivatives”. Additionally, the “Alkaloids and derivatives”, “Organic acids and derivatives” and “Organic nitrogen compounds” appear less well explained by MetFrag (perhaps with more rearrangements, or less distinguishable spectra), such that the improvement from the RO approach is more apparent. For SIRIUS, “Lipids and lipid-like molecules” as well as “Organic oxygen compounds” benefit the most from LC-MS^2^Struct in top-1 (both improving by 5.7 percentage units) and top-20 (4.1 and 3.2 percentage units, respectively). In general, for “Lipid and lipid-like molecules”, LC-MS^2^Struct seems to achieve the best improvement (top-1 and top-20) over all MS^2^ scorers. However, depending on the MS^2^ scorer this improvement distributes differently across the lipid sub-classes (see Extended Data Figure 2), such as “Fatty Acyls”, “Prenol lipids” or “Sphingolipids”.

**Figure 3:**
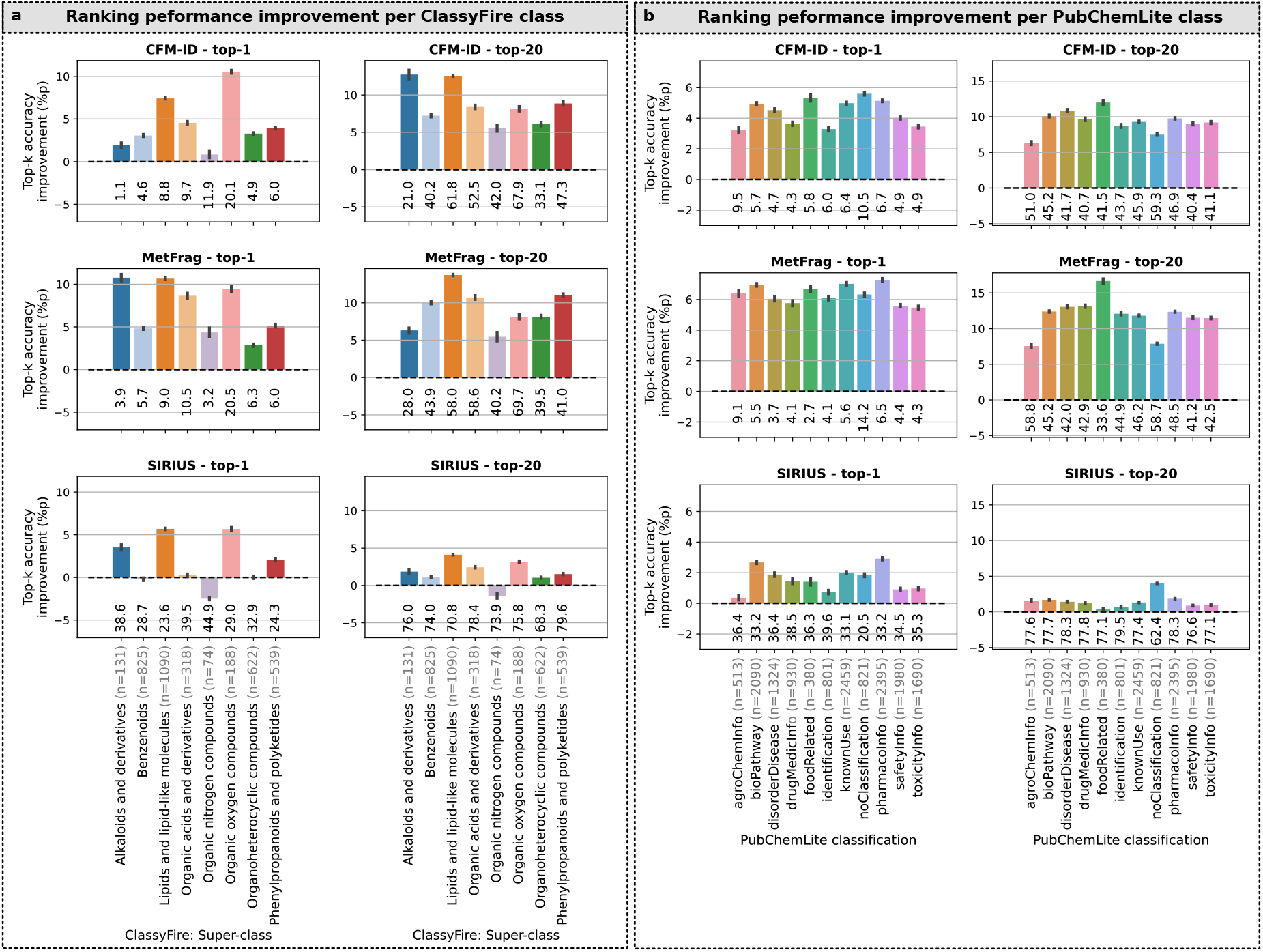
Performance gain by LC-MS^2^Struct across molecular classes. The figure shows the average (50 samples) and 95%-confidence interval (1000 bootstrapping samples) of the ranking performance (top-*k*) improvement of LC-MS^2^Struct compared to Only-MS^2^ (baseline). The top-*k* accuracies (%) under the bars show the Only-MS^2^ performance. For each molecular class, the number of unique molecular structures in the class is denoted in the x-axis label (n). **a**: Molecular classification using the ClassyFire [51] framework (Class level). **b**: PubChemLite [40] annotation classification system. Molecules not present in PubChemLite are summarized under the “noClassification” category. Note that in PubChemLite a molecule can belong to multiple categories.

For the PubChemLite classification (Figure 3**b**) we also see that the MS^2^ scorers benefit differently from LC-MS^2^Struct. The improvement is generally close to the average improvement of the respective MS^2^ scorers and seems more equally distributed across the annotation categories.

For example for CFM-ID, the biggest top-1 improvements are in the “Food Related” and “noClassification” categories. On the other hand, for SIRIUS the “PharmacoInfo” and “BioPathway” categories improve the most. MetFrag shows the most consistent performance improvement across the categories. “AgroChemInfo” benefits the least from LC-MS^2^Struct (top-1 and top-20) A possible explanation could be that the molecules categorized as agrochemicals are mainly “Benzenoids” (48.5%), “Organoheterocyclic compounds” (25.9%) and “Organic acids and derivatives” (11.6%). As shown in Figure 3**a**, these three ClassyFire classes show low (CFM-ID and MetFrag) or practically no (SIRIUS) improvement when using ROs.

### Annotation of stereoisomers

Finally, we study whether LC-MS^2^Struct can annotate stereoisomers more accurately than MS^2^ alone, considering differences between stereoisomers that vary in their double-bond orientation (*e*.*g. cis-trans* or *E-Z* isomerism), which may potentially lead to differences in their LC-behavior (see Figure 5**a**). We consider candidate sets containing stereoisomers and evaluate LC-MS^2^Struct only using MassBank records where the ground truth structure has stereochemistry information provided, *i*.*e*. where the InChIKey second block is not “UHFFFAOYSA” (ONLYSTEREO data setup, see Methods). The molecular candidates are represented using two different molecular fingerprints: One that includes stereochemistry information (3D); and one that omits it (2D) (see Methods). This allows us to assess the importance of stereochemistry-aware features for the structure annotation.

Figure 4**a** shows the ranking performance of LC-MS^2^Struct using 2D and 3D fingerprints. When looking into the top-1 performance of LC-MS^2^Struct (3D) for the individual MS^2^ scorers, we observe an improvement by 2.6, 3.8 and 3.2 percentage units for CFM-ID, MetFrag and SIRIUS, respectively. This translates to performance gains of 87.3%, 95.9% and 44.3%.

**Figure 4:**
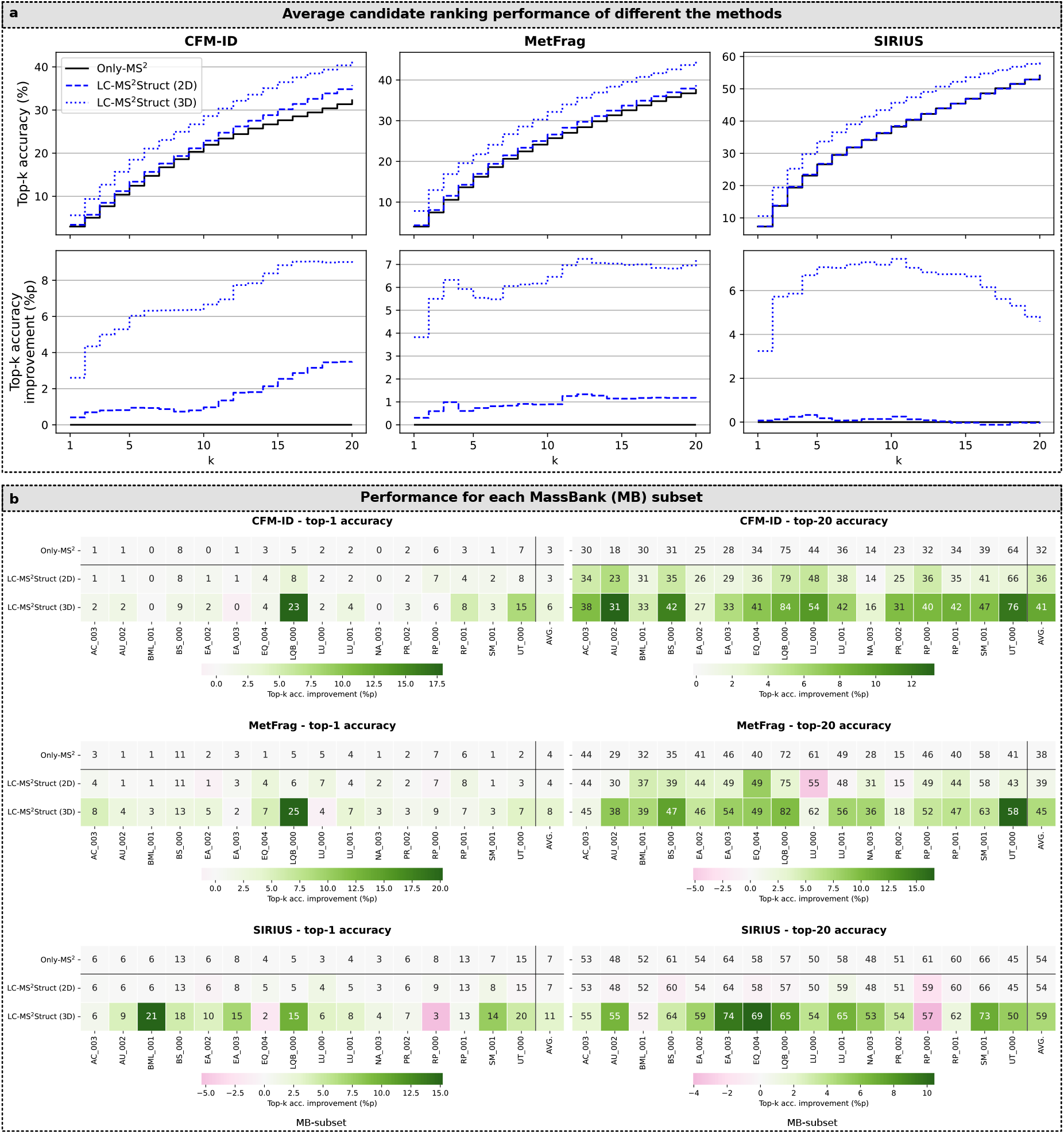
Using LC-MS^2^Struct to identify stereoisomers. **a**: Comparison of the performance, measured by top-*k* accuracy, of LC-MS^2^Struct using either 2D (no stereochemistry) or 3D (with stereochemistry) molecular fingerprints in the ONLYSTEREO setting. The results shown are averaged accuracies over 94 sample MS feature sequences (LC-MS^2^ experiments). **b**: Average top-*k* accuracies per MassBank (MB) subset rounded to full integers. The color encodes the performance improvement of each score integration method compared with Only-MS^2^.

Generally, LC-MS^2^Struct improves the ranking for all three MS^2^ scorers. The improvement, however, is notably larger when using stereochemistry-aware (3D) candidate features. Interestingly, a similar behavior could be observed in the ALLDATA setting (see Extended Data Figure 3), even though the absolute performance improvements were smaller. This experiment demonstrates that LC-MS^2^Struct can use RO information to improve the annotation of stereoisomers.

## Discussion

LC-MS^2^Struct is a novel approach for the integration of tandem mass spectrometry (MS^2^) and liquid chromatography (LC) data for the structural annotation of small molecules. The method learns from the pairwise dependencies in the retention order (RO) of MS features within similar LC configurations and can generalize across different, heterogeneous LC configurations. Furthermore, the use of stereochemistry-aware molecular fingerprints enables LC-MS^2^Struct to annotate stereoisomers in LC-MS^2^ experiments based on the observed ROs. Also, our novel processing pipeline to group all (MS^2^, RT)-data from MassBank into subsets of homogeneous LC-MS^2^ conditions, which is implemented and made available in the “massbank2db” [52] Python package will, we believe, make MassBank more accessible to other researchers and hence lower the bar of entry to computational metabolomics research.

Our experiments demonstrate that LC-MS^2^Struct annotates small molecules with an accuracy far superior to more traditional retention time (RT) filtering and log*P*-based approaches, and also markedly better than previous methods that rely on ROs. In particular, compared to Bach et al. [34], who used a graphical model as a post-hoc integration tool of MS^2^ scores and RO predictions, the benefits of learning the parameters of the graphical model are clear. All three studied MS^2^ scorers could be improved by LC-MS^2^Struct, including the best-of-class SIRIUS, for which improvements have generally been hard to come by due to its already high baseline accuracy. Our results show the superiority of stereochemistry-aware molecular features for the structure annotation of LC-MS^2^ data. Remarkably, this was not only the case for the annotation of stereoisomers but also for candidates only distinguished by their 2D structure. This result could be relevant for improving structural annotations in ion mobility separation-mass spectrometry (IMS-MS) with collision cross section (CCS) measurements.

Our examples indicated that LC-MS^2^Struct separates candidates with varying double-bond stereochemistry, *i*.*e. E/Z-* and *cis/trans*-isomers (see *e*.*g*. Figure 5). However, there were very few examples of double-bond and/or chiral isomers measured on the same LC system in our dataset, which makes it difficult to quantify this effect, or interrogate these further until more such data is publicly available. Furthermore, as non-chiral LC cannot distinguish stereoisomers that only differ in their chiral centers, the development of more selective stereochemistry-aware molecular features, ignoring the chiral annotations, might be beneficial. We also note that the direct modelling of a node score (MS^2^ information) predictor in the Structured Support Vector Machine would be possible. However, as the MS^2^ scorers used here are already relatively mature and well-known in the community, we have left this research line open for future efforts.

**Figure 5:**
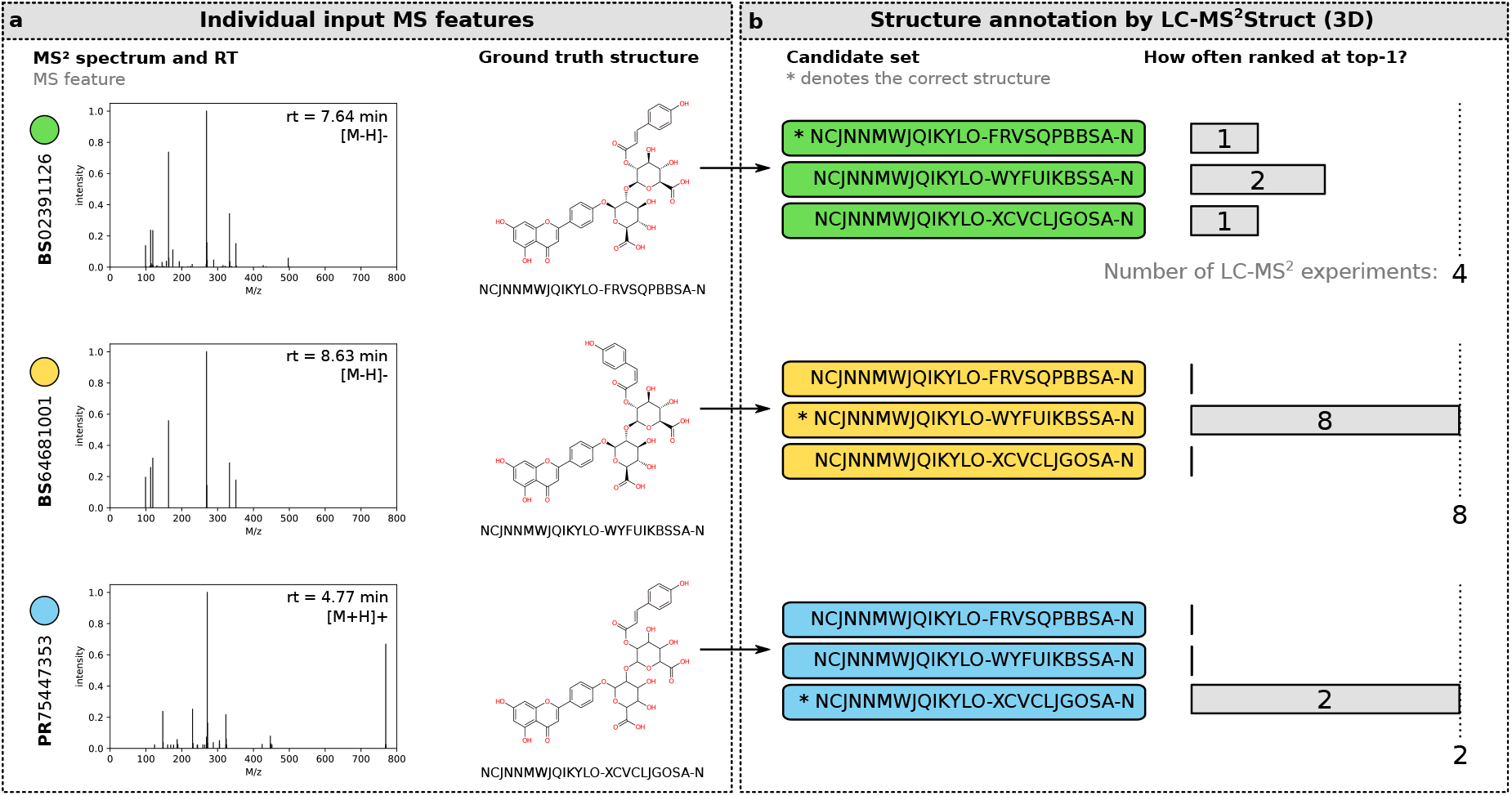
Application of LC-MS^2^Struct to annotate stereoisomers. Post-hoc analysis of the stereoisomer annotation using LC-MS^2^Struct for three (MS^2^, RT)-tuples from our MassBank data associated with the same 2D skeleton (InChIKey first block). In our evaluation, all three MS features were analysed multiple times in different contexts (BS02391126 in 4, BS64681001 in 8 and PR75447353 in 2 LC-MS^2^ experiments). **a**: MS features with their ground truth annotations. Two of the spectra (starting with BS) were measured under the same LC condition (MB-subset “BS_000”), demonstrating the separation of *E/Z-*isomers on LC columns. **b**: The candidate sets of the three features are identical (defined by the molecular formula C_36_H_32_O_19_) and only contain three structures. For 12 out of the 14 LC-MS^2^ experiments, LC-MS^2^Struct predicts the correct *E/Z-*isomer.

## Methods

### Notation

We use the following notation to describe LC-MS^2^Struct:

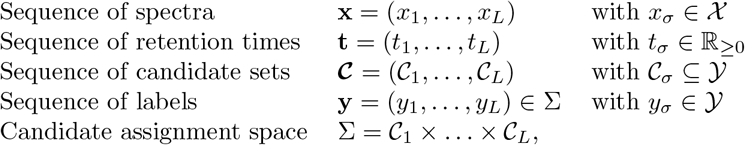

where 𝒳 and 𝒴 denote the MS^2^ spectra and the molecular structure space, respectively, and 𝒞 denotes a candidate set that is a subset of all possible molecular structures, and *A* × *B* denotes cross product of two sets *A* and *B*. For the purpose of model training and evaluation, we assume a dataset with ground truth labeled MS feature sequences: 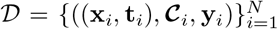, where *N* denotes the total number of sequences. We use *i, j* ∈ ℕ_≥0_ to index MS feature sequences and *σ, τ* ∈ ℕ_≥0_ as indices for individual MS features within a sequence, e.g. *x*_*iσ*_ denotes the MS^2^ spectrum at index *σ* in the sequence *i*. The length of a sequence of MS features is denoted with *L*. We denote the ground truth labels (candidate assignment) of sequence *i* with **y**_*i*_ and any labelling with **y**. Both, **y**_*i*_ and **y** are in Σ_*i*_. We use *y* to denote the candidate label variable, whereas *m* denotes a particular molecular structure. For example, *y*_*σ*_ = *m* means, that we assign the molecular structure *m* as label to the MS feature *σ*.

### Graphical model for joint annotation of MS features

We consider the molecular annotation problem for the output of an LC-MS^2^, that means assigning a molecular structure to each MS feature, as a structured prediction problem [35, 47, 46], relying on a graphical model representation of the sets of MS features arising from an LC-MS^2^ experiment. For each MS feature *σ* we want to predict a label *y*_*σ*_ from a fixed and finite candidate (label) set 𝒞_*σ*_. We model the observed retention orders (RO) between each MS feature pair (*σ, τ*) within an LC-MS^2^ experiment, as pairwise dependencies of the features. We define an undirected graph *G* = (*V, E*) with the vertex set *V* containing a node *σ* for each MS feature and the edge set *E* containing an edge for each MS feature pair *E* = {(*σ, τ*) | *σ, τ* ∈ *V, σ* ≠ *τ*} (c.f. Figure 1**a** and **c**). The resulting graph is complete with an edge between all pairs of nodes. This allows us to make use of arbitrary pairwise dependencies, instead of limiting to, say, adjacent retention times. This modeling choice was previously shown to be beneficial by Bach et al. [34]. Here we extend that approach by learning from the pairwise dependencies to optimize joint annotation accuracy, which leads to markedly improved annotation accuracy.

For learning, we define a scoring function *F* that, given the input MS feature sequences (**x, t**) and its corresponding sequence of candidate sets **𝒞**, computes a compatibility score between the measured data and *any* possible sequence of labels **y** ∈ Σ:

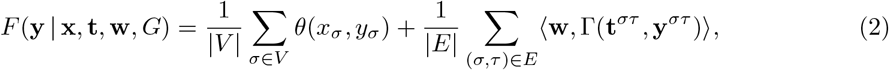

where *θ* : 𝒳 × 𝒴 → (0, 1] is a function returning an MS^2^ matching score between the spectrum *x*_*σ*_ and a candidate *y*_*σ*_ ∈ 𝒞_*σ*_, ⟨·, ·⟩ denotes the inner product, and **w** is a model weight vector to predict the RO matching score, based on the joint feature vector Γ : ℝ_≥0_ × ℝ_≥0_ × 𝒴 × 𝒴 → ℱ between the observed RO derived from **t**^*στ*^ = (*t*_*σ*_, *t*_*τ*_) and a pair of molecular candidates **y**^*στ*^ = (*y*_*σ*_, *y*_*τ*_).

Equation (2) consists of two parts: (1) A score computed over the nodes in *G* capturing the MS^2^ information; and (2) a score expressing the agreement of observed and predicted RO computed over the edge set. We assume that the node scores are pre-computed by a MS^2^ scorer such as CFM-ID [10], MetFrag [14] or SIRIUS [9]. The node scores are normalized to (0, 1] within each candidate set _*σ*_. The edge scores are predicted for each edge (*σ, τ*) using the model **w** and the joint-feature vector Γ:

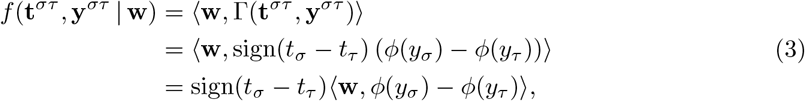

with *ϕ* : 𝒴 → ℱ _𝒴_ being a function embedding a molecular structure into a feature space. The edge prediction function (3) will produce a height edge score, if the observed RO (*i*.*e*. sign(*t*_*σ*_ − *t*_*τ*_)) agrees with the predicted one.

Using the compatibility score function (2), the predicted joint annotation for (**x, t**) corresponds to the the highest scoring label sequence 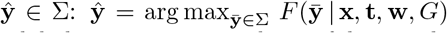. In practice, however, instead of only predicting the best label sequence, it can be useful to *rank* the molecular candidates *m* ∈ 𝒞_*σ*_ for each MS feature *σ*. That is because for state-of-the-art MS^2^ scorers, the annotation accuracy in the top-20 candidate list is typically much higher than for the highest ranked candidate (top-1).

Our framework provides candidate rankings by solving the following problem for each MS feature *σ* and *m* ∈ 𝒞_*σ*_:

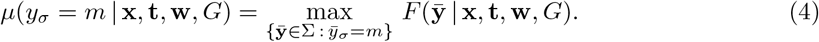

Problem (4) returns a max-marginal *μ* score for each candidate *m*. That is, the maximum compatibility score *any* label sequence 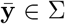 with 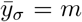 can achieve. One can interpret Equation (2) as the log-space representation of a unnormalized Markov Random Field probability distribution over **y** associated with an undirected graphical model *G* [44].

### Feasible inference using random spanning trees (RST)

For general graphs *G*, the maximum a posterior (MAP) inference problem (*i*.*e*. finding the highest scoring label sequence **y** given an MS feature sequence) is an 𝒩𝒫-hard problem [53, 54]. The max-marginals inference (MMAP), needed for the candidate ranking, is an even harder problem which is 𝒩𝒫^PP^ complete [54]. However, efficient inference approaches have been developed. In particular, if *G* is tree-like, we can efficiently compute the max-marginals using dynamic programming and the max-product algorithm [43, 44]. Such tree-based approximations have shown to be successful in various practical applications [45, 46, 34].

Here, we follow the work by Bach et al. [34] and sample a set of random spanning trees (RST) 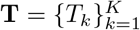 from *G*, whereby *K* denotes the size of the RST sample. Each tree *T*_*k*_ has the same node set *V* as *G*, but and an edge set *E*(*T*) ⊆ *E*, with |*E*(*T*)| = *L* − 1, ensuring that *T* is a single connected component and cycle free. We follow the sampling procedure used by Bach et al. [34]. Given the RST set **T** we compute the averaged max-marginals to rank the molecular candidates [34]:

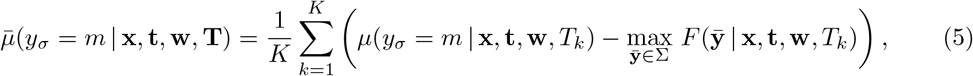

where we subtract the maximum compatibility score from the marginal values corresponding to the individual trees to normalize the marginals before averaging [34]. This normalization value can be efficiently computed given the max-marginals *μ*. In our experiments, we train *K* individual models (**w**_*k*_) and associate them with the trees *T*_*k*_ to increase the diversity. The influence of the number of SSVM models on the prediction performance is shown in Extended Data Figure 4.

### The Structured Support Vector Machine (SSVM) model

To train the model parameters **w** (see Equation (2)), we implemented a variant of the Structured Support Vector Machine (SSVM) [36, 35]. Its primal optimization problem is given as [55]:

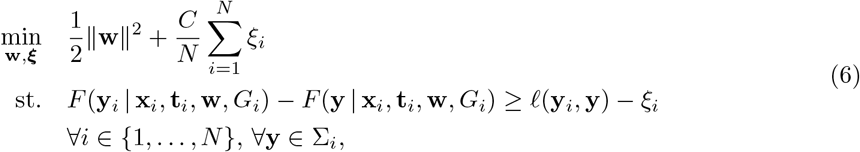

where *C >* 0 being the regularization parameter, *ξ*_*i*_ ≥ 0 is the slack variable for example *i* and 𝓁 : Σ_*i*_ × Σ_*i*_ → ℝ_≥0_ being a function capturing the loss between two label sequences. The constraint set definition (st.) of problem (6) leads to a parameter vector **w** that is trained according to the max-margin principle [36, 35, 47], that is the score *F* (**y**_*i*_) of the correct label should be greater than the score *F* (**y**) of any other label sequence by at least the specified margin 𝓁(**y**_*i*_, **y**). Note that in the SSVM problem (6) a different graph *G*_*i*_ = (*V*_*i*_, *E*_*i*_) can be associated to each training example *i*, allowing, for example, to process sequences of different length.

We solve (6) in its dual formulation and use the Frank-Wolfe algorithm [56] following the recent work by Lacoste-Julien et al. [55]. In the supplementary material we derive the dual problem and demonstrate how to solve it efficiently using the Frank-Wolfe algorithm and RST approximations for *G*_*i*_. Optimizing the dual problem enables us to use non-linear kernel functions *λ* : 𝒴 × 𝒴 → ℝ_≥0_ measuring the similarity between the molecular structures associated with the label sequences.

The label loss function 𝓁 is defined as follows:

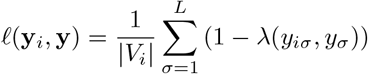

and satisfies 𝓁(**y, y**) = 0 (a required property [55]), if *λ* is a normalized kernel, which holds true in our experiments (we used the MinMax kernel [57]).

### Pre-processing pipeline for raw MassBank records

Extended Data Figure 5 illustrates our MassBank (MB) pre-processing pipeline implemented in the Python package “massbank2db” [52]. First, the MassBank records text files were parsed and the MS^2^ spectrum, ground truth annotation, RT and meta-information extracted. Records with missing MS^2^, RT or annotation were discarded. We use the MB 2020.11 release for our experiments.

Subsequently, we grouped the MassBank records into subsets (denoted as MB-subsets) where the (MS^2^, RT)-tuples were measured under the same LC- and MS-conditions.

Supplementary Table 1 summarizes the grouping criteria. In the next step, we used the InChIKey [58] identifier in MassBank to retrieve the SMILES [59] representation from PubChem [20] (1 Feb., 2021), rather than using the contributor-supplied SMILES. This ensures a consistent SMILES source for the molecular candidates and ground truth annotations.

Three more filtering steps were performed before creating the final database, to remove records: (1) if the ground truth exact mass deviated too far (*>* 20ppm) from the calculated exact mass based on the precursor mass-per-charge and adduct type; (2) if the subset contained *<* 50 unique molecular structures; (3) if they were potential isobars (see pull-request #152 in the MassBank GitHub repository, https://github.com/MassBank/MassBank-data/pull/152).

Supplementary Table 3 summarizes the LC-MS^2^ meta-information for all generated MB-subsets.

### Generating the molecular candidate sets

We used SIRIUS [11, 9] to generate the molecular candidate sets. For each MassBank record the ground truth molecular formula was used by SIRIUS to collect the candidate structures from PubChem [20]. The candidate sets generated by SIRIUS contain a single stereoisomer per candidate, identified by their InChIKey first block (structural skeleton). To study the ability of LC-MS^2^Struct to annotate the stereochemical variant of the molecules, we enriched the SIRIUS candidates sets with stereoisomers, using the InChIKey first block of each candidate to search PubChem (1 Feb., 2021) for stereoisomers. The additional molecules were then added to the candidate sets.

### Pre-computing the MS^2^ matching scores

For each MB-subset, MS^2^ spectra with identical adduct type (e.g. `[M+H]+`) and ground truth molecular structure were aggregated. Depending on the MS^2^ scorer, we either merged the MS^2^ into a single spectrum (CFM-ID and MetFrag) following the strategy by Ruttkies et al. [14] or we provided the MS^2^ spectra separately (SIRIUS). For the spectra merging we used the “mzClust hclust” function of the xcms package [60], which first combines all MS^2^ spectra’s peaks into a single peaklist and subsequently merges peaks based on a mass error threshold.

To compute the CFM-ID (v4.0.7) MS^2^ matching score, we first predicted the in silico MS^2^ spectra for all molecular candidate structures based on their isomeric SMILES representation using the pre-trained CFM-ID models (Metlin 2019 MSML) by Wang et al. [10]. We merged the three in silico spectra predicted by CFM-ID for different collision energies and compared them with the merged MassBank spectrum using the modified cosine similarity [61] implemented in the matchms [62] (v0.9.2) Python library. For MetFrag (v2.4.5), the MS^2^ matching scores were calculated using the `FragmenterScore` feature based on the isomeric SMILES representation of the candidates. For SIRIUS, the required fragmentation trees are computed using the ground truth molecular formula of each MassBank spectrum. SIRIUS uses canonical SMILES and hence does not encode stereochemical information (which is absent in the canonical SMILES). Therefore, we used the same SIRIUS MS^2^ matching score for all stereoisomers sharing the same InChIKey first block.

For all three MS^2^ scorers we normalized the MS^2^ matching scores to the range [0, 1] separately for each candidate set. For the machine learning based scorers (CFM-ID and SIRIUS), the matching scores of the candidates associated with a particular MassBank record using in evaluation were predicted using models that did include its ground truth structure (determined by InChIKey first block).

If a MS^2^ scorer failed on a MassBank record, we assigned a constant MS^2^ score to each candidate.

### Molecular feature representations

Extended connectivity fingerprints with function-classes (FCFP) [38] were used to represent molecular structures in our experiments. We employed RDKit (v2021.03.1) to generate *counting* FCFP fingerprints. The fingerprints were computed based on the isomeric SMILES, using the parameter “useChirality” to generate fingerprints that either encoded stereochemistry (3D) or not (2D). To define the set of substructures in the fingerprint vector, we first generated all possible substructures, using a FCFP radius of two, based on a set of 50000 randomly sampled molecular candidates associated with our training data, and all the ground truth training structures, resulting in 6925 (3D) and 6236 (2D) substructures. We used 3D FCFP fingerprints in our experiments, except for the experiments focusing on the annotation of stereoisomers, where we used both 2D and 3D fingerprints for comparison. We used the MinMax-kernel [57] to compute the similarity between the molecules.

### Computing molecular categories

For the analysis of the ranking performance for different molecular categories, we used two classification systems, ClassyFire [51], which classifies molecules according to their structure and PubChemLite [40], which classifies molecules according to information available for 10 exposomics-relevant categories. For ClassyFire, we used the “classyfireR” R package to retrieve the classification for each ground truth molecular structure in our dataset. For PubChemLite, the classification categories were retrieved via InChIKey first block matching of each molecular structure; if it was not found in PubChemLite, the category “noClassification” was assigned.

### Training and evaluation data setups

We only considered MassBank data that has been analyzed using a LC reversed phase (RP) column. We removed molecules from the data if their measured retention time (RT) was less than three times the estimated column dead-time [63], as we considered such molecules to be non-retaining.

We considered two separate data setups. The first one, denoted by ALLDATA, used all available MassBank data to train and evaluate LC-MS^2^Struct. This setup was used to compare the different candidate ranking approaches as well as to investigate the performance across various molecular classes. The second setup, denoted by ONLYSTEREO, used MassBank records where the ground truth molecular structure contains stereochemical information, *i*.*e*. where the InChIKey second block is not “UHFFFAOYSA”. This setup was used in the experiments regarding the ability of LC-MS^2^Struct to distinguish stereochemistry. In the training, we additionally used MassBank records that appear only without stereochemical information in our candidate sets, identified by the InChIKey second block equal to “UHFFFAOYSA” in PubChem. The number of available training and evaluation (MS^2^, RT)-tuples per MB-subset are summarized in Supplementary Table 2.

For each MB-subset we sampled a set of LC-MS^2^ experiments, i.e. (MS^2^, RT)-tuple sequences, from the available evaluation data. The number of LC-MS^2^ experiments (*n* below) depended on the number of available (MS^2^, RT)-tuples (see Supplementary Table 2) as follows:

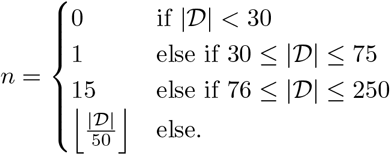

where 𝒟 is a set of (MS^2^, RT)-tuples with ground truth annotation and molecular candidate sets associated with a MB-subset. If there are less than 30 (MS^2^, RT)-tuples available, we do not generate an evaluation LC-MS^2^ experiment from the corresponding MB-subset. Based on this sampling scheme, we obtained 354 and 94 LC-MS^2^ experiments for ALLDATA and ONLYSTEREO, respectively, for our evaluation (see Supplementary Table 2).

We trained eight (*K* = 8) separate SSVM models **w**_*k*_ for each evaluation LC-MS^2^ experiment. For each SSVM model we first generated a set containing the (MS^2^, RT)-tuples from all MB-subsets. Then, we removed all tuples whose ground truth molecular structure, determined by the InChIKey first block, was in the respective evaluation LC-MS^2^ experiment. Lastly, we randomly sampled LC-MS^2^ experiments from the training tuples, within their respective MB-subset, with a length randomly chosen from 4 to (maximum) 32 (see also Figure 1**e**) and an RST *T*_*ik*_ assigned for each MS feature sequence *i*. In total 768 LC-MS^2^ training experiments were generated for each SSVM model. To speed up the model training, we restricted the candidate set size |𝒞_*iσ*_| of each training MS feature *σ* to maximum 75 candidate structures by random sub-sampling. We ensure that the correct candidate is included in the sub-sample. Each SSVM model **w**_*k*_ was applied to the evaluation LC-MS^2^ experiment, associated with different RSTs *T*_*k*_, and the averaged max-marginal scores where used for the final candidate ranking (see Equation (5) and Figure 1**c**).

### SSVM hyper-parameter optimization

The SSVM regularization parameter *C* was optimized for each training set separately using grid search and evaluation on a random validation set sampled from the training data’s (MS^2^, RT)-tuples (33%). A set of LC-MS^2^ experiments was generated from the validation set and used to determine the Normalized Discounted Cumulative Gain (NDCG) [64] for each *C* value. The regularization parameter with the highest NDCG value was chosen to train the final model. We used the scikit-learn [65] (v0.24.1) Python package to compute the NDCG value, taking into account ranks up until 10 (NDCG@10) and defined the relevance for each candidate to be 1 if it is the correct one and 0 otherwise. To reduce the training time, we searched the optimal *C*^*^ only for SSVM model *k* = 0 and used *C*^*^ for the other models with *k >* 0.

### Ranking performance evaluation

We computed the ranking performance (top-*k* accuracy) for a given LC-MS^2^ experiment using the tie-breaking strategy described in [11]: If a ranking method assigns an identical score to a set of *n* molecular candidates, then all accuracies at the ordinal ranks *k* at which one of these candidates is found are increased by 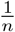. We computed a candidate score (*i*.*e*. Only-MS^2^, LC-MS^2^Struct, etc.) for each molecular structure in the candidate set (identified by PubChem CID). Depending on the data setup (see Supplementary Table 4) we first collapse the candidates by InChIKey first block (ALLDATA, method comparison and molecule category analysis) or full InChIKey (ONLYSTEREO stereochemistry prediction), assigning the maximum candidate score for each InChIKey first block or InChIKey group, respectively. Subsequently, we compute the top-*k* accuracy based on the collapsed candidate sets.

For the performance analysis of individual molecule categories, either ClassyFire [51] or PubChemLite [40] classes, we first computed the rank of the correct molecular structure for each (MS^2^, RT)-tuple of each LC-MS^2^ evaluation experiment based on Only-MS^2^ and LC-MS^2^Struct scores. Subsequently, we computed the top-*k* accuracy for each molecule category, associated with at least 50 unique ground truth molecular structures (based on InChIKey first block). As a ground truth structure can appear multiple times in our dataset, we generate 50 random samples, each containing only one example per unique structure, and computed the averaged top-*k* accuracy.

### Comparison of LC-MS^2^Struct with other approaches

We compared LC-MS^2^Struct with three different approaches to integrate tandem mass spectrum (MS^2^) and retention time (RT) information, namely RT filtering, log*P* prediction and retention order (RO) prediction.

For RT filtering (MS^2^+RT), we followed Aicheler et al. [26] who used the relative error 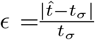, between the predicted 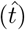 and observed (*t*_*σ*_) retention time. We set the filtering threshold to the 95%-quantile of the relative RT prediction errors estimated from the RT model’s training data, following [27, 29]. We used scikit-learn’s [65] (v0.24.1) implementation of the Support Vector Regression (SVR) [66] with radial basis function (RBF) kernel for the RT prediction. For SVR, we use the same 196 features, computed using RDKit (v2021.03.1), as Bouwmeester et al. [25].

For log*P* prediction (MS^2^+log*P*) we followed Ruttkies et al. [14] who assigned a weighted sum of an MS and log*P* score 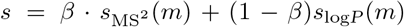 to each candidate *m* ∈ 𝒞_*σ*_, and use it rank the set of molecular candidates. The log*P* score is given by *s*_log*P*_(*m*) = 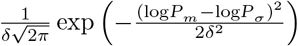, where log*P*_*m*_ is the predicted XLogP3 [50] extracted from PubChem [20] for candidate *m*, and log*P*_*σ*_ = *a* · *t*_*σ*_ + *b* is the XLogP3 value of the unknown compound, associated with MS feature *σ*, predicted based on its measured RT *t*_*σ*_. The parameters *a* and *b* of the linear regression model were determined using a set of RT and XLogP3 tuples associated with the LC system. As Ruttkies et al. [14], we set the *δ* = 1.5 and set *β* such that it optimizes the top-1 candidate ranking accuracy, calculated from a set of 25 randomly generated training LC-MS^2^ experiments.

For retention order prediction (MS^2^+RO) we used the approach by Bach et al. [34] which relies on a Ranking Support Vector Machine (RankSVM) implementation in the Python library ROSVM [31, 67] (v0.5.0). We used counting `substructure` fingerprints calculated using CDK (v2.5) [68] and the MinMax kernel [57]. The MS^2^ matching scores and predicted ROs were used to compute max-marginal ranking scores using the framework by Bach et al. [34]. We used the author’s implementation in version 0.2.3 [69]. The hyper-parameters *β* and *k* of the model were optimized for each evaluation LC-MS^2^ experiment separately using the respective training data. To estimate *β* we generated 25 LC-MS^2^ experiments from the training data and selected the *β* that maximized the Top20AUC [34] ranking performance. The sigmoid parameter *k* was estimated using Platt’s method [70] calibrated using RankSVM’s training data. We used 128 random spanning trees per evaluation LC-MS^2^ experiment to compute the averaged max-marginals.

For the experiments comparing the different methods we used all LC-MS^2^ experiments generated, except the ones from the MB-subsets “CE_001”, “ET_002”, “KW_000” and “RP_000” (see Supplementary Table 2). For those subsets the evaluation LC-MS^2^ experiment contain all available (MS^2^, RT)-tuples, leaving no LC system specific data to train the RT (MS^2^+RT) or log*P* (MS^2^+log*P*) prediction models. The RT and log*P* prediction models are trained in a structure disjoint fashion using the RT data of the particular MB-subset associated with the evaluation LC-MS^2^. The RO prediction model used by MS^2^+RO is trained structure disjoint as well, but using the RTs of all MB-subsets.

## Supporting information

MassBank subsets meta data

## Data availability

All data used in our experiments is available online [71] (https://zenodo.org/record/5854661). The candidate rankings of all LC-MS^2^ experiments are available online: ALLDATA [72] (https://zenodo.org/record/6451016) and ONLYSTEREO [73] (https://zenodo.org/record/6037629).

## Code availability

The source code developed for this study is available on GitHub, including the implementation of LC-MS^2^Struct [74] (v2.13.0, https://github.com/aalto-ics-kepaco/msms_rt_ssvm); scripts to run the experiments [75] (https://github.com/aalto-ics-kepaco/lcms2struct_exp); and the library implementing the MassBank pre-processing [52] (v0.9.0, https://github.com/bachi55/massbank2db). The candidate fingerprints were computed by the ROSVM Python library [67] (v0.5.0, https://github.com/bachi55/rosvm) using RDKit (2021.03.1). The SSVM library uses the max-marginal inference solver implemented by Bach et al. [34] (v0.2.3, https://github.com/aalto-ics-kepaco/msms_rt_score_integration).

## Acknowledgments

The work by E.B. and J.R. was partially supported by Academy of Finland grants 310107 (MA-COME) and 334790 (MAGITICS). E.L.S. acknowledges funding support from the Luxembourg National Research Fund (FNR) for project A18/BM/12341006 and discussions with Drs. Greg Landrum (ETHZ) and Evan Bolton (NCBI/NLM/NIH). The authors acknowledge CSC-IT Center for Science, Finland and Aalto Science-IT infrastructure, Finland for generous computational resources. E.B. thanks Dr. K. Dührkop for generating the SIRIUS candidate sets and predicting the SIRIUS MS^2^ scores.

## Authors contributions

E.B. and J.R. designed the research. E.B. implemented the MassBank pre-processing. E.B. developed, implemented and evaluated the computational method. E.B., E.L.S. and J.R. interpreted the results. E.B., E.L.S. and J.R. wrote the manuscript.

## Extended data figures

**Extended Data Figure 1:**
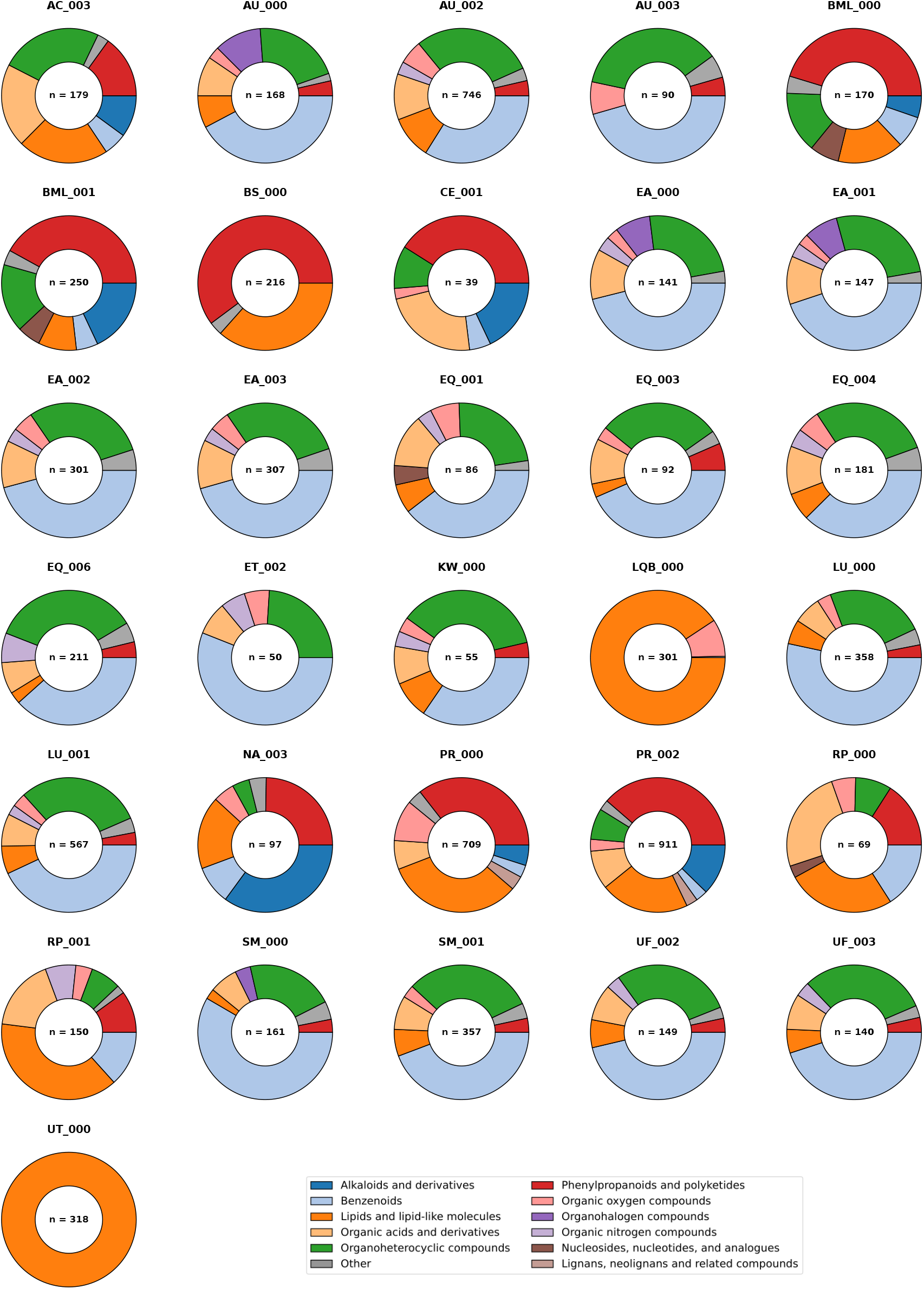
Distribution of molecule classes in the MassBank (MB) subsets. Classy-Fire SuperClass distribution [51] for each MB-subset studied in our experiments. Within each MB-subset, the label “Other” is assigned to each SuperClass which makes up less then 2.5% of all molecules. The center label represents the number of examples for the respective MB-subset.

**Extended Data Figure 2:**
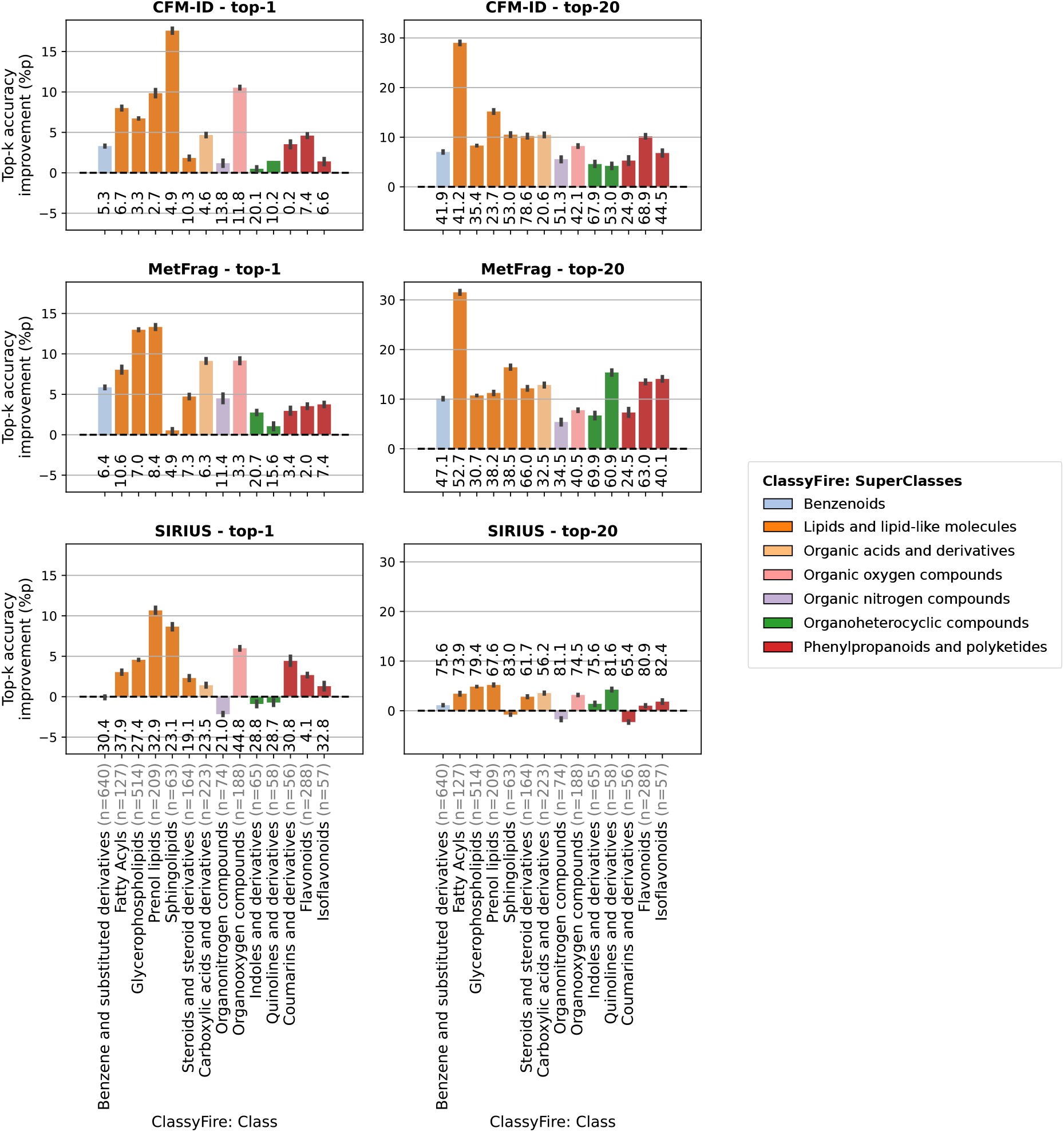
Performance gain by LC-MS^2^Struct across ClassyFire Class-level annotations. The Classes (shown in the bars) are colour coded by SuperClasses (see legend). The figure shows the average (50 samples) and 95%-confidence interval (1000 bootstrapping samples) of the ranking performance (top-*k*) improvement of LC-MS^2^Struct compared to Only-MS^2^ (baseline). The top-*k* accuracies (%) under the bars show the Only-MS^2^ performance. For each molecule class, the number of unique molecular structures in the class is denoted in the x-axis label (n).

**Extended Data Figure 3:**
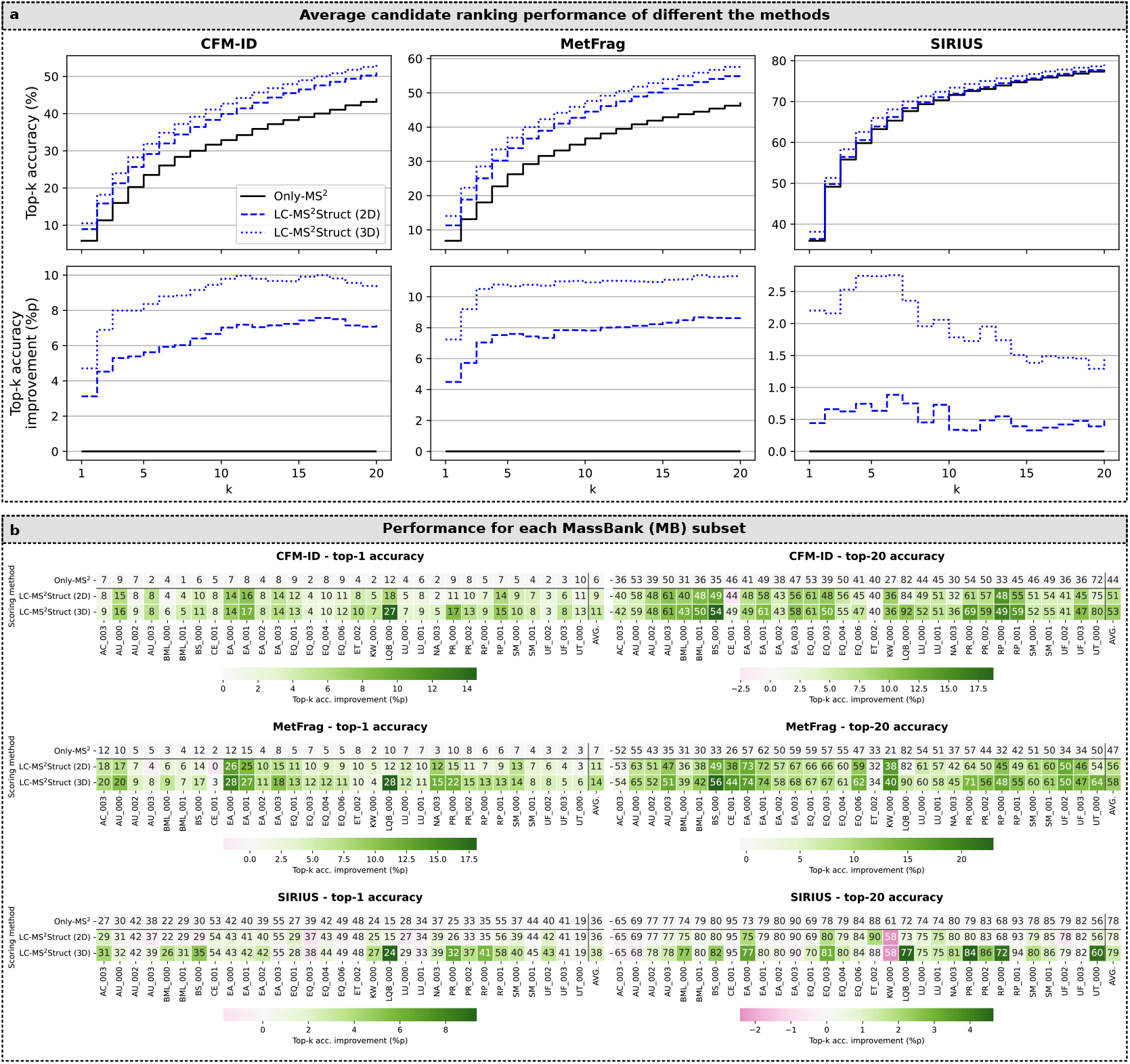
Using LC-MS^2^Struct with different molecule feature representations to identify the correct structure at the level of first InChIKey block (InChIKey-1). **a**: Comparison of the performance, measured by top-*k* accuracy, of LC-MS^2^Struct using either 2D (no stereochemistry) or 3D (with stereochemistry) molecular fingerprints in the ALLDATA setting. The results shown are averaged accuracies over 354 sample MS feature sequences (LC-MS^2^ experiments). **b**: Average top-*k* accuracies per MassBank (MB) subset rounded to full integers. The color encodes the performance improvement of each score integration method compared to Only-MS^2^.

**Extended Data Figure 4:**
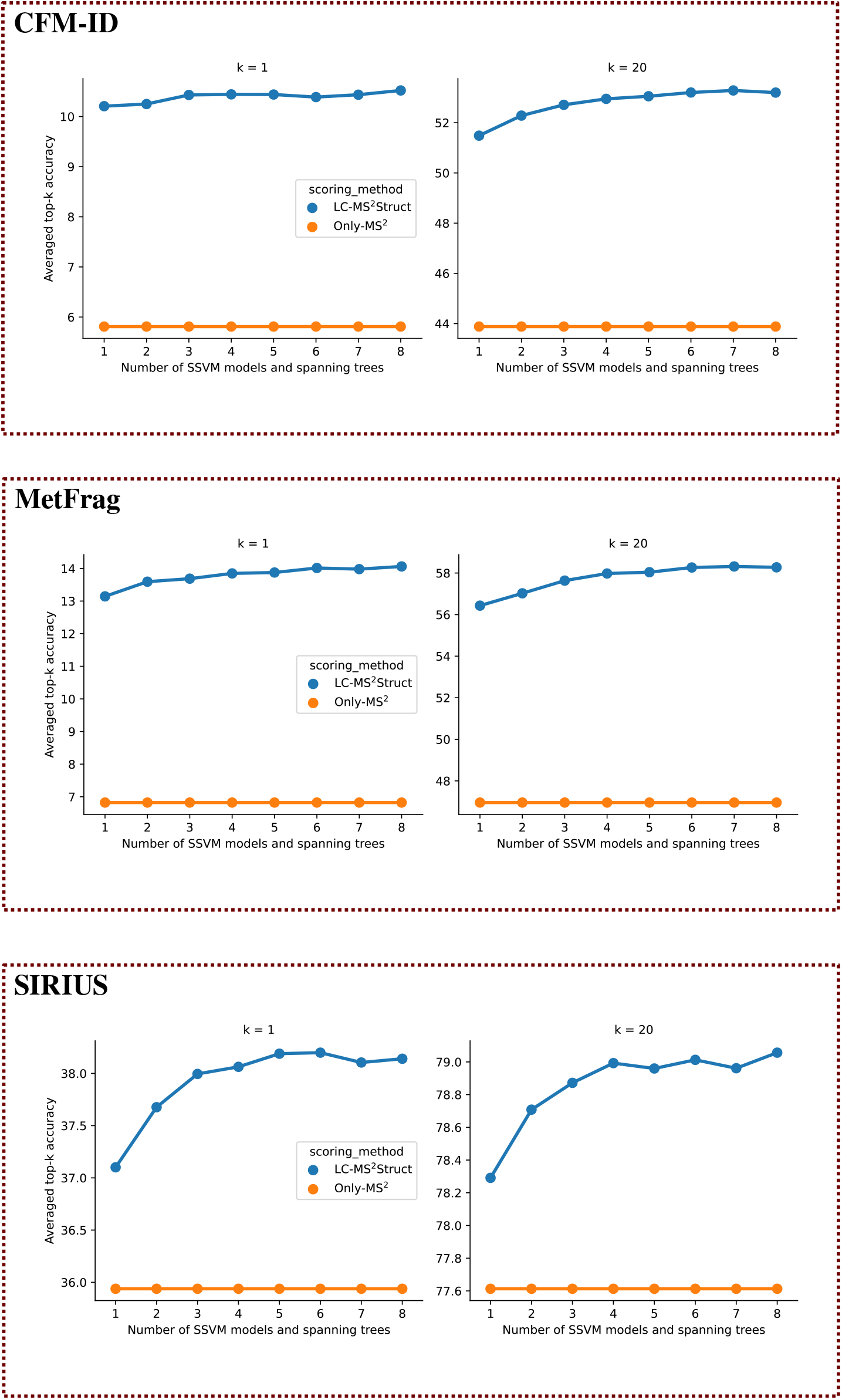
Performance comparison of LC-MS^2^Struct against using only MS^2^ information (Only-MS^2^) for different number of SSVM models. The performance curves for the three MS^2^-scorers are shown separately. The top-*k* accuracies shown are averaged accuracies over 354 sample MS feature sequences (LC-MS^2^ experiments) from the ALLDATA setting.

**Extended Data Figure 5:**
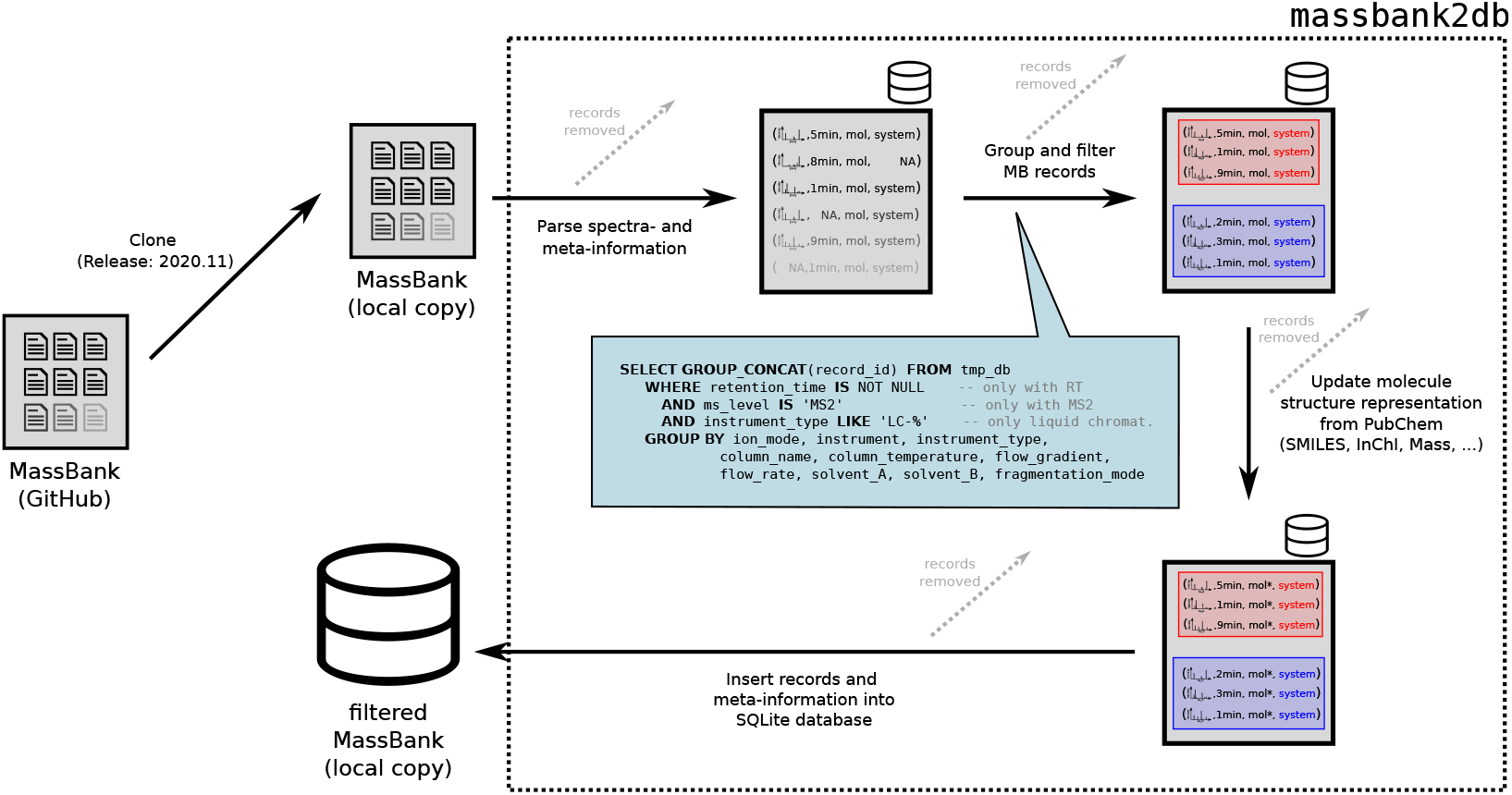
Processing pipeline of the MassBank data. Illustration of the processing pipeline to extract the training data from MassBank. The depicted workflow is implemented in the “massbank2db” Python package [52].

