## Supplementary information for "Joint structural annotation of small molecules using liquid chromatography retention order and tandem mass spectrometry data"

### 1 Structured Support Vector Machine (SSVM)

In this section we first derive dual formulation of the primal Structured Support Vector Machine (SSVM) problem presented in the main manuscript (see Methods "The Structured Support Vector Machine (SSVM) model"). Subsequently we show how the SSVM optimization problem is solved in the dual space.

#### 1.1 SSVM: From primal to dual problem

We introduce the following notation:

$$\Delta\boldsymbol{\theta}_i(\mathbf{y}) = \frac{1}{|V_i|}\sum_{\sigma\in V_i}\left(\theta(x_{i\sigma}, y_{i\sigma}) - \theta(x_{i\sigma}, y_\sigma)\right) \qquad \text{MS}^2 \text{ score difference}$$

$$\langle \mathbf{w}, \Delta\boldsymbol{\Gamma}_i(\mathbf{y})\rangle = \frac{1}{|E_i|}\sum_{(\sigma,\tau)\in E_i}\langle \mathbf{w}, \Gamma(\mathbf{t}_i^{\sigma\tau}, \mathbf{y}_i^{\sigma\tau}) - \Gamma(\mathbf{t}_i^{\sigma\tau}, \mathbf{y}^{\sigma\tau})\rangle \qquad \text{Retention order score difference}$$

$$\ell_i(\mathbf{y}) = \ell(\mathbf{y}_i, \mathbf{y}) = \frac{1}{|V_i|}\sum_{\sigma\in V_i}\left(1 - \lambda(y_{i\sigma}, y_\sigma)\right) \qquad \text{Label losses}$$

using $\langle \mathbf{a}, \mathbf{b} \rangle = \mathbf{a}^T\mathbf{b}$ to express the inner product between two vectors $\mathbf{a}, \mathbf{b} \in \mathbb{R}^d$ and re-formulate the primal SSVM using it:

$$\begin{aligned} \min_{\mathbf{w},\boldsymbol{\xi}} \quad & \frac{1}{2}\|\mathbf{w}\|^2 + \frac{C}{N}\langle \mathbf{1}, \boldsymbol{\xi} \rangle \\ \text{st.} \quad & \left( \frac{\ell_i(\mathbf{y}) - \Delta\boldsymbol{\theta}_i(\mathbf{y})}{|V_i|} - \frac{\langle \mathbf{w}, \Delta\boldsymbol{\Gamma}_i(\mathbf{y}) \rangle}{|E_i|} \right) - \xi_i \leq 0 \\ & \forall i \in \{1, \dots, N\}, \forall \mathbf{y} \in \Sigma_i, \end{aligned}$$

with $\boldsymbol{\xi} \in \mathbb{R}^N_{\geq 0}$ is a vector storing all slack-variables $\xi_i$. Note that we have the ground truth label $\mathbf{y}_i$ in the label space $\Sigma_i$ and therefore implicitly include $-\xi_i \leq 0 \Rightarrow \xi_i \geq 0$ in the set of constraints.

We can now set up the Lagrangian $\mathcal{L}$, with a dual variable $\alpha(i, \mathbf{y}) \geq 0$ for all example $i$ and label sequence $\mathbf{y} \in \Sigma_i$:

$$\begin{aligned} \mathcal{L}(\mathbf{w}, \boldsymbol{\xi}, \boldsymbol{\alpha}) &= \frac{1}{2}\mathbf{w}^T\mathbf{w} + \frac{C}{N}\boldsymbol{\xi}^T\mathbf{1} + \sum_{i=1}^{N}\sum_{\mathbf{y}\in\Sigma_i} \alpha(i,\mathbf{y}) \left( \frac{\ell_i(\mathbf{y}) - \Delta\boldsymbol{\theta}_i(\mathbf{y})}{|V_i|} - \frac{\mathbf{w}^T\Delta\boldsymbol{\Gamma}_i(\mathbf{y})}{|E_i|} - \xi_i \right) \\ &= \frac{1}{2}\mathbf{w}^T\mathbf{w} + \frac{C}{N}\boldsymbol{\xi}^T\mathbf{1} + \sum_{(i,\mathbf{y})} \alpha(i,\mathbf{y}) \frac{\ell_i(\mathbf{y}) - \Delta\boldsymbol{\theta}_i(\mathbf{y})}{|V_i|} - \sum_{(i,\mathbf{y})} \alpha(i,\mathbf{y}) \frac{\mathbf{w}^T\Delta\boldsymbol{\Gamma}_i(\mathbf{y})}{|E_i|} - \dots \\ &\quad \sum_{i=1}^{N} \xi_i \sum_{\mathbf{y}\in\Sigma_i} \alpha(i,\mathbf{y}) \\ &= \frac{1}{2}\mathbf{w}^T\mathbf{w} + \boldsymbol{\xi}^T \left( \frac{C}{N}\mathbf{1} - \mathbf{B}\mathbf{1} \right) + \sum_{(i,\mathbf{y})} \alpha(i,\mathbf{y}) \frac{\ell_i(\mathbf{y}) - \Delta\boldsymbol{\theta}_i(\mathbf{y})}{|V_i|} - \sum_{(i,\mathbf{y})} \alpha(i,\mathbf{y}) \frac{\mathbf{w}^T\Delta\boldsymbol{\Gamma}_i(\mathbf{y})}{|E_i|}, \end{aligned} \tag{1}$$

where $\mathbf{B} \in \mathbb{R}^{N\times M}_{\geq 0}$ ($M = \sum_i |\Sigma_i|$, total number of *all* possible label sequences) is a matrix storing all dual variables $\alpha(i, \mathbf{y})$ with the following structure:

$$\mathbf{B} = \begin{bmatrix} \boldsymbol{\alpha}_1^T & 0 & 0 & \dots & 0 \\ 0 & \boldsymbol{\alpha}_2^T & 0 & \dots & 0 \\ & & \vdots & & \\ 0 & 0 & \dots & 0 & \boldsymbol{\alpha}_N^T \end{bmatrix},$$

with $\boldsymbol{\alpha}_i = (\alpha(i, \mathbf{y}))_{\mathbf{y}\in\Sigma_i} \in \mathbb{R}^{M_i}_{\geq 0}$.

We now differentiate $\mathcal{L}$ with respect to $\mathbf{w}$ and $\boldsymbol{\xi}$

$$\frac{\partial \mathcal{L}(\mathbf{w}, \boldsymbol{\xi}, \boldsymbol{\alpha})}{\partial \mathbf{w}} = \mathbf{w} - \sum_{(i,\mathbf{y})} \alpha(i,\mathbf{y}) \frac{\Delta\boldsymbol{\Gamma}_i(\mathbf{y})}{L_i} \stackrel{!}{=} 0 \Rightarrow \boxed{\mathbf{w}^* = \sum_{(i,\mathbf{y})} \alpha(i,\mathbf{y}) \frac{\Delta\boldsymbol{\Gamma}_i(\mathbf{y})}{|E_i|}} \tag{2}$$

$$\frac{\partial \mathcal{L}(\mathbf{w}, \boldsymbol{\xi}, \boldsymbol{\alpha})}{\partial \boldsymbol{\xi}} = \frac{C}{N}\mathbf{1} - \mathbf{B}\mathbf{1} \stackrel{!}{=} 0 \Rightarrow \boxed{\sum_{\mathbf{y}\in\Sigma_i} \alpha(i,\mathbf{y}) = \frac{C}{N} \quad \forall i \in \{1, \dots, N\}}. \tag{3}$$

and plug the solutions (2) and (3) back to $\mathcal{L}$ (Equation (1)):

$$g(\boldsymbol{\alpha}) = \mathcal{L}(\mathbf{w}^*, \boldsymbol{\xi}^*, \boldsymbol{\alpha}) = -\frac{1}{2}\sum_{(i,\mathbf{y})}\sum_{(j,\bar{\mathbf{y}})} \alpha(i,\mathbf{y})\alpha(j,\bar{\mathbf{y}}) \frac{\langle \Delta\boldsymbol{\Gamma}_i(\mathbf{y}), \Delta\boldsymbol{\Gamma}_j(\bar{\mathbf{y}}) \rangle}{|E_i| \cdot |E_j|} + \sum_{(i,\mathbf{y})} \alpha(i,\mathbf{y}) \frac{\ell_i(\mathbf{y}) - \Delta\boldsymbol{\theta}_i(\mathbf{y})}{|V_i|} \tag{4}$$

to arrive at the dual optimization problem:

$$\begin{aligned} \max_{\boldsymbol{\alpha}} \quad & -\frac{1}{2}\sum_{(i,\mathbf{y})}\sum_{(j,\bar{\mathbf{y}})} \alpha(i,\mathbf{y})\alpha(j,\bar{\mathbf{y}}) \frac{\langle \Delta\boldsymbol{\Gamma}_i(\mathbf{y}), \Delta\boldsymbol{\Gamma}_j(\bar{\mathbf{y}}) \rangle}{|E_i| \cdot |E_j|} + \sum_{(i,\mathbf{y})} \alpha(i,\mathbf{y}) \frac{\ell_i(\mathbf{y}) - \Delta\boldsymbol{\theta}_i(\mathbf{y})}{|V_i|} \\ \text{st.} \quad & \sum_{\mathbf{y}\in\Sigma_i} \alpha(i,\mathbf{y}) = \frac{C}{N} \quad \forall i \in \{1, \dots, N\} \\ & \alpha(i,\mathbf{y}) \geq 0 \quad \forall i, \forall \mathbf{y} \in \Sigma_i. \end{aligned} \tag{5}$$

**Algorithm 1** Mini-Batch Conditional Gradient Algorithm to solve Problem (5) respectively (6) [1]

$k \leftarrow 0$
Let $\boldsymbol{\alpha}^{(k)} \in \mathcal{A}$ ▷ A feasible initial dual variable
**for** $e \in \{1, \ldots, n_{\text{epochs}}\}$ **do**
  $\mathcal{I} \leftarrow$ `split_into_random_batches`$(\{1, \ldots, N\})$
  **for** $\mathcal{I}_B \in \mathcal{I}$ **do**
    $\mathbf{s} \leftarrow \boldsymbol{\alpha}^{(k)}$
    **for** $i \in \mathcal{I}_B$ **do** ▷ Sub-problem
      Compute $\hat{\mathbf{y}}_i = \arg\max_{\mathbf{y} \in \Sigma_i} \frac{\ell(\mathbf{y}_i, \mathbf{y}) + \boldsymbol{\theta}(\mathbf{x}_i, \mathbf{y})}{|V_i|} + \frac{\langle \boldsymbol{\Gamma}(\mathbf{t}_i, \mathbf{y}), \mathbf{A}\boldsymbol{\alpha}^{(k)} \rangle}{|E_i|}$
      $\mathbf{s}_i \leftarrow \left( 0, \ldots, 0, \underbrace{\frac{C}{N}}_{\text{at index for: } \hat{\mathbf{y}}_i}, 0, \ldots, 0 \right)$ ▷ Update Direction
    **end for**
    $\gamma \leftarrow$ `determine_step_size_using_linesearch()` ▷ Line-search. Fixed rule possible, e.g., $\frac{2N}{k+2N}$
    $\boldsymbol{\alpha}^{(k+1)} \leftarrow \boldsymbol{\alpha}^{(k)} + \gamma(\mathbf{s} - \boldsymbol{\alpha}^{(k)})$ ▷ Update
    $k \leftarrow k + 1$
  **end for**
**end for**

To simplify the representation of the dual problem, we define the following notation:

$$\boldsymbol{\alpha}_i = (\alpha(i, \mathbf{y}))_{\mathbf{y} \in \Sigma_i} \in \mathbb{R}_{\geq 0}^{M_i} \qquad \text{Dual variables for example } i$$

$$\boldsymbol{\alpha} = (\boldsymbol{\alpha}_i)_{i=1}^{N} \in \mathbb{R}_{\geq 0}^{M} \qquad \text{Dual variables}$$

$$\mathbf{A} = \left\{ \frac{\Delta \boldsymbol{\Gamma}_i(\mathbf{y})}{|E_i|} \mid i \in \{1, \ldots, N\}, \mathbf{y} \in \Sigma_i \right\} \in \mathbb{R}^{|\mathcal{F}| \times M} \qquad \text{Edge-feature differences}$$

$$\mathbf{b} = \left( \frac{\ell_i(\mathbf{y}) - \Delta \boldsymbol{\theta}_i(\mathbf{y})}{|V_i|} \right)_{i \in \{1, \ldots, N\}, \mathbf{y} \in \Sigma_i} \in \mathbb{R}^{M} \qquad \text{Label-losses \& MS}^2 \text{ score differences}$$

$$\mathcal{A}_i = \left\{ \boldsymbol{\alpha}_i \in \mathbb{R}^{M_i} \mid \alpha(i, \mathbf{y}) \geq 0 \ \forall \mathbf{y} \in \Sigma_i, \sum_{\mathbf{y} \in \Sigma_i} \alpha(i, \mathbf{y}) = \frac{C}{N} \right\} \qquad \text{Feasible space of } \boldsymbol{\alpha}_i$$

$$\mathcal{A} = \bigcup_{i=1}^{N} \mathcal{A}_i \qquad \text{Feasible space of } \boldsymbol{\alpha}$$

Using this notation, we can re-write Problem (5) into:

$$\max_{\boldsymbol{\alpha} \in \mathcal{A}} \quad g(\alpha) = -\frac{1}{2} \langle \mathbf{A}\boldsymbol{\alpha}, \mathbf{A}\boldsymbol{\alpha} \rangle + \langle \boldsymbol{\alpha}, \mathbf{b} \rangle. \tag{6}$$

#### 1.2 Solving the dual SSVM optimization problem

We use the Frank-Wolfe algorithm [2] (also known as Conditional Gradient Method, see [3, 4, 1] for more recent literature on the topic) to solve the SSVM in its dual representation (6). This approach to solve the SSVM has been previously proposed in the literature [5, 1]. We in particular follow the idea of [1] and apply a block-coordinate variant of the original Frank-Wolfe algorithm. However, instead of updating only a single variable $i$ per iteration we update a subset, or *mini-batch*, $\mathcal{I}_B$ of the variables. Our Frank-Wolfe implementation is outlined in Algorithm 1. In the following we will explain its main steps.

##### 1.2.1 Solving the sub-problem to find the update direction

In each iteration of Algorithm 1 we need to solve the so called the *loss-augmented decoding* problem [1], which essentially means we need to find the highest scoring label sequence $\hat{\mathbf{y}}_i$ (see Methods "Label sequence prediction using graph inference"). Using $\hat{\mathbf{y}}_i$ we can define a feasible update direction for the gradient descent.

**Partial derivative of $g(\boldsymbol{\alpha})$:** Given equation (4) we can derive its partial derivative:

$$\underset{i}{\nabla} g(\boldsymbol{\alpha}) = \mathbf{b}_i - \langle \mathbf{A}_i, \mathbf{A}\boldsymbol{\alpha} \rangle \in \mathbb{R}^{M_i}$$

with $\mathbf{A}_i = \left\{ \frac{\Delta \boldsymbol{\Gamma}_i(\mathbf{y})}{|E_i|} \,|\, \mathbf{y} \in \Sigma_i \right\} \in \mathbb{R}^{|\mathcal{F}| \times M_i}$, i.e. the columns of $\mathbf{A}$ belonging to example $i$, and similarly $\mathbf{b}_i = \left( \frac{\ell_i(\mathbf{y}) - \Delta \boldsymbol{\theta}_i(\mathbf{y})}{|V_i|} \right)_{\mathbf{y} \in \Sigma_i} \in \mathbb{R}^{M_i}$.

**Update direction of the Frank-Wolfe algorithm:** Note, as we strive to maximize the dual objective (Equation (6)) we need to find an update direction $\mathbf{s}_i \in \mathcal{A}_i$ that is the solution of the following problem:

$$\begin{aligned}
\mathbf{s}_i &= \underset{\mathbf{s}'_i \in \mathcal{A}_i}{\arg\max} \left\langle \underset{i}{\nabla} g(\boldsymbol{\alpha}), \left( \mathbf{s}'_i - \boldsymbol{\alpha}_i^{(k)} \right) \right\rangle \\
&= \underset{\mathbf{s}'_i \in \mathcal{A}_i}{\arg\max} \left\langle \underset{i}{\nabla} g(\boldsymbol{\alpha}), \mathbf{s}'_i \right\rangle - \underbrace{\left\langle \underset{i}{\nabla} g(\boldsymbol{\alpha}), \boldsymbol{\alpha}_i^{(k)} \right\rangle}_{\text{constant}} \\
&= \underset{\mathbf{s}'_i \in \mathcal{A}_i}{\arg\max} \left\langle \underset{i}{\nabla} g(\boldsymbol{\alpha}), \mathbf{s}'_i \right\rangle
\end{aligned} \tag{7}$$

The solution of equation (7) is defined over the simplex $\mathcal{A}_i$. Therefore, it is sufficient to find the coordinate of $\underset{i}{\nabla} g(\boldsymbol{\alpha})$ with the maximum (gradient) value and assign $\frac{C}{N}$ to the corresponding index in $\mathbf{s}_i$ [3]. Note that each coordinate corresponds to a label sequence $\mathbf{y}$ in the label space $\Sigma_i$ of example $i$. With that in mind, we can rewrite the equation (7) as follows:

$$\begin{aligned}
\hat{\mathbf{y}}_i &= \underset{\mathbf{y} \in \Sigma_i}{\arg\max} \; \underset{i}{\nabla} g(\boldsymbol{\alpha}) \\
&= \underset{\mathbf{y} \in \Sigma_i}{\arg\max} \; \frac{\ell_i(\mathbf{y}) - \Delta \boldsymbol{\theta}_i(\mathbf{y})}{|V_i|} - \left\langle \frac{\Delta \boldsymbol{\Gamma}_i(\mathbf{y})}{|E_i|}, \mathbf{A}\boldsymbol{\alpha} \right\rangle \\
&= \underset{\mathbf{y} \in \Sigma_i}{\arg\max} \; \frac{\ell(\mathbf{y}_i, \mathbf{y}) - (\boldsymbol{\theta}(\mathbf{x}_i, \mathbf{y}_i) - \boldsymbol{\theta}(\mathbf{x}_i, \mathbf{y}))}{|V_i|} - \left\langle \frac{\boldsymbol{\Gamma}(\mathbf{t}_i, \mathbf{y}_i) - \boldsymbol{\Gamma}(\mathbf{t}_i, \mathbf{y})}{|E_i|}, \mathbf{A}\boldsymbol{\alpha} \right\rangle \\
&= \underset{\mathbf{y} \in \Sigma_i}{\arg\max} \; \frac{\ell(\mathbf{y}_i, \mathbf{y}) - \boldsymbol{\theta}(\mathbf{x}_i, \mathbf{y}_i) + \boldsymbol{\theta}(\mathbf{x}_i, \mathbf{y})}{|V_i|} - \left\langle \frac{\boldsymbol{\Gamma}(\mathbf{t}_i, \mathbf{y}_i)}{|E_i|}, \mathbf{A}\boldsymbol{\alpha} \right\rangle + \left\langle \frac{\boldsymbol{\Gamma}(\mathbf{t}_i, \mathbf{y})}{|E_i|}, \mathbf{A}\boldsymbol{\alpha} \right\rangle \\
&= \underset{\mathbf{y} \in \Sigma_i}{\arg\max} \; \frac{\ell(\mathbf{y}_i, \mathbf{y}) + \boldsymbol{\theta}(\mathbf{x}_i, \mathbf{y})}{|V_i|} + \left\langle \frac{\boldsymbol{\Gamma}(\mathbf{t}_i, \mathbf{y})}{|E_i|}, \mathbf{A}\boldsymbol{\alpha} \right\rangle - \underbrace{\left( \frac{\boldsymbol{\theta}(\mathbf{x}_i, \mathbf{y}_i)}{|V_i|} + \left\langle \frac{\boldsymbol{\Gamma}(\mathbf{t}_i, \mathbf{y}_i)}{|E_i|}, \mathbf{A}\boldsymbol{\alpha} \right\rangle \right)}_{\text{Constant for all } \mathbf{y}} \\
&= \underset{\mathbf{y} \in \Sigma_i}{\arg\max} \; \underbrace{\frac{\ell(\mathbf{y}_i, \mathbf{y}) + \boldsymbol{\theta}(\mathbf{x}_i, \mathbf{y})}{|V_i|}}_{\text{Node-scores}} + \underbrace{\frac{\langle \boldsymbol{\Gamma}(\mathbf{t}_i, \mathbf{y}), \mathbf{A}\boldsymbol{\alpha} \rangle}{|E_i|}}_{\text{Edge-scores}} \\
&= \underset{\mathbf{y} \in \Sigma_i}{\arg\max} \; \frac{\ell(\mathbf{y}_i, \mathbf{y})}{|V_i|} + F(\mathbf{y} \,|\, \mathbf{x}_i, \mathbf{t}_i, \mathbf{A}\boldsymbol{\alpha}, G_i),
\end{aligned} \tag{8}$$

where we use the definition of the scoring function $F$ (see Methods "Graphical model for joint annotation of MS features.") and the relationship between the primal and dual parameters, i.e. $\mathbf{w} = \mathbf{A}\boldsymbol{\alpha}$. In our experiments we use a random spanning tree (RST) $T_i$ to approximate the full graph $G_i$. We sample a RST for each example $i$ in our training data. Using RST to approximate the graphical model enables efficient inference of the loss-augmented decoding problem (8) [6] (see Methods "Feasible inference using random spanning trees (RST)").

##### 1.2.2 Stepsize determination using line-search

We follow [1] and derive the optimal step-size $\gamma_{\text{LS}}$ using the linear-search approach. For that, we first need to solve the following optimization problem:

$$\begin{aligned}\gamma_{\text{opt}} &= \arg\max_{\gamma\in[0,1]} g(\boldsymbol{\alpha}+\gamma(\mathbf{s}-\boldsymbol{\alpha}))\\ &= \arg\max_{\gamma\in[0,1]} -\frac{1}{2}\langle \mathbf{A}(\boldsymbol{\alpha}+\gamma(\mathbf{s}-\boldsymbol{\alpha})), \mathbf{A}(\boldsymbol{\alpha}+\gamma(\mathbf{s}-\boldsymbol{\alpha}))\rangle + \langle \boldsymbol{\alpha}+\gamma(\mathbf{s}-\boldsymbol{\alpha}), \mathbf{b}\rangle.\end{aligned} \tag{9}$$

Generally, problem (9) can be solved by setting its derivative, an univariate quadratic function of $\gamma$, to zero:

$$\begin{aligned}\frac{\partial g}{\partial \gamma} &= \langle \mathbf{b}, \mathbf{s}-\boldsymbol{\alpha}\rangle - \langle \mathbf{A}(\boldsymbol{\alpha}+\gamma(\mathbf{s}-\boldsymbol{\alpha})), \mathbf{A}(\mathbf{s}-\boldsymbol{\alpha})\rangle \overset{!}{=} 0\\ &\Leftrightarrow \langle \mathbf{b}, \mathbf{s}-\boldsymbol{\alpha}\rangle = \langle \mathbf{A}(\boldsymbol{\alpha}+\gamma(\mathbf{s}-\boldsymbol{\alpha})), \mathbf{A}(\mathbf{s}-\boldsymbol{\alpha})\rangle\\ &\Leftrightarrow \langle \mathbf{b}, \mathbf{s}-\boldsymbol{\alpha}\rangle = \langle \mathbf{A}\boldsymbol{\alpha}, \mathbf{A}(\mathbf{s}-\boldsymbol{\alpha})\rangle + \gamma\langle \mathbf{A}(\mathbf{s}-\boldsymbol{\alpha}), \mathbf{A}(\mathbf{s}-\boldsymbol{\alpha})\rangle\\ &\Leftrightarrow \langle \mathbf{b}, \mathbf{s}-\boldsymbol{\alpha}\rangle - \langle \mathbf{A}\boldsymbol{\alpha}, \mathbf{A}(\mathbf{s}-\boldsymbol{\alpha})\rangle = \gamma\langle \mathbf{A}(\mathbf{s}-\boldsymbol{\alpha}), \mathbf{A}(\mathbf{s}-\boldsymbol{\alpha})\rangle\\ &\Leftrightarrow \gamma_{\text{opt}} = \frac{\langle \mathbf{b}, \mathbf{s}-\boldsymbol{\alpha}\rangle - \langle \mathbf{A}\boldsymbol{\alpha}, \mathbf{A}(\mathbf{s}-\boldsymbol{\alpha})\rangle}{\langle \mathbf{A}(\mathbf{s}-\boldsymbol{\alpha}), \mathbf{A}(\mathbf{s}-\boldsymbol{\alpha})\rangle}\\ &\Leftrightarrow \gamma_{\text{opt}} = \frac{\langle\langle \mathbf{b} - \langle \mathbf{A}, \mathbf{A}\boldsymbol{\alpha}\rangle, \mathbf{s}-\boldsymbol{\alpha}\rangle}{\langle \mathbf{A}(\mathbf{s}-\boldsymbol{\alpha}), \mathbf{A}(\mathbf{s}-\boldsymbol{\alpha})\rangle}\\ &\Leftrightarrow \boxed{\gamma_{\text{opt}} = \frac{\langle \nabla g(\boldsymbol{\alpha}), \mathbf{s}-\boldsymbol{\alpha}\rangle}{\langle \mathbf{A}(\mathbf{s}-\boldsymbol{\alpha}), \mathbf{A}(\mathbf{s}-\boldsymbol{\alpha})\rangle}}\end{aligned} \tag{10}$$

After computing the optimal step-size $\gamma_{\text{opt}}$ using equation (10), we derive the line-search step-size by restricting $\gamma_{\text{opt}}$ to $[0, 1]$:

$$\gamma_{\text{LS}} = \max(0, \min(1, \gamma_{\text{opt}})).$$

When training the SSVM using the stochastic block coordinate descent outlined in Algorithm 1, we can exploit the fact that the difference vector $(\mathbf{s}-\boldsymbol{\alpha})$ is zero except for the $i$'th block (respectively all blocks $i \in \mathcal{I}_{\mathcal{B}}$). In the following the we derive the nominator and denominator of equation (10) needed to compute the step-size in practice.

**Nominator of equation (10):**

$$\begin{aligned}&\langle \nabla g(\boldsymbol{\alpha}), (\mathbf{s}-\boldsymbol{\alpha})\rangle\\ &\quad= \sum_{i\in\mathcal{I}_{\mathcal{B}}}\sum_{\mathbf{y}\in\Sigma_i}(s(i,\mathbf{y})-\alpha(i,\mathbf{y}))\left(\frac{\ell_i(\mathbf{y})-\Delta\boldsymbol{\theta}_i(\mathbf{y})}{|V_i|} - \left\langle \frac{\Delta\boldsymbol{\Gamma}_i(\mathbf{y})}{|E_i|}, \mathbf{A}\boldsymbol{\alpha}\right\rangle\right)\\ &\quad= \sum_{i\in\mathcal{I}_{\mathcal{B}}}\sum_{\mathbf{y}\in\Sigma_i}(s(i,\mathbf{y})-\alpha(i,\mathbf{y}))\left(\frac{\ell(\mathbf{y}_i,\mathbf{y})+\boldsymbol{\theta}(\mathbf{x}_i,\mathbf{y})}{|V_i|} + \left\langle \frac{\boldsymbol{\Gamma}(\mathbf{t}_i,\mathbf{y})}{|E_i|}, \mathbf{A}\boldsymbol{\alpha}\right\rangle - \dots\right.\\ &\qquad\qquad \left.\left(\frac{\boldsymbol{\theta}(\mathbf{x}_i,\mathbf{y}_i)}{|V_i|} + \left\langle \frac{\boldsymbol{\Gamma}(\mathbf{t}_i,\mathbf{y}_i)}{|E_i|}, \mathbf{A}\boldsymbol{\alpha}\right\rangle\right)\right)\\ &\quad= \sum_{i\in\mathcal{I}_{\mathcal{B}}}\sum_{\mathbf{y}\in\Sigma_i}(s(i,\mathbf{y})-\alpha(i,\mathbf{y}))\left(\underbrace{\frac{\ell(\mathbf{y}_i,\mathbf{y})}{|V_i|} + F(\mathbf{y}\,|\,\mathbf{x}_i,\mathbf{t}_i,\mathbf{A}\boldsymbol{\alpha},G_i)}_{\text{Compare with equation. (8)}} - F(\mathbf{y}_i\,|\,\mathbf{x}_i,\mathbf{t}_i,\mathbf{A}\boldsymbol{\alpha},G_i)\right)\end{aligned}$$

$$= \sum_{i \in \mathcal{I}_\mathcal{B}} \sum_{\mathbf{y} \in \Sigma_i} (s(i,\mathbf{y}) - \alpha(i,\mathbf{y})) \left( \frac{\ell(\mathbf{y}_i, \mathbf{y})}{|V_i|} + F(\mathbf{y} \,|\, \mathbf{x}_i, \mathbf{t}_i, \mathbf{A}\boldsymbol{\alpha}, G_i) \right) - \ldots$$

$$\sum_{i \in \mathcal{I}_\mathcal{B}} F(\mathbf{y}_i \,|\, , \mathbf{x}_i, \mathbf{t}_i, \mathbf{A}\boldsymbol{\alpha}, G_i) \underbrace{\sum_{\mathbf{y} \in \Sigma_i} (s(i,\mathbf{y}) - \alpha(i,\mathbf{y}))}_{=0 \text{ (Definition of feasible set)}}$$

$$= \sum_{i \in \mathcal{I}_\mathcal{B}} \sum_{\mathbf{y} \in \Sigma_i} (s(i,\mathbf{y}) - \alpha(i,\mathbf{y})) \left( \underbrace{\frac{\ell(\mathbf{y}_i, \mathbf{y})}{|V_i|} + F(\mathbf{y} \,|\, , \mathbf{x}_i, \mathbf{t}_i, \mathbf{A}\boldsymbol{\alpha}, G_i)}_{\equiv \mathbf{I}} \right),$$

with $\boldsymbol{\theta}(\mathbf{x}_i, \mathbf{y}) = \sum_{\sigma=1}^{L_i} \theta(x_{i\sigma}, y_\sigma)$.

**Denominator of equation** (10)**:**

$$\langle \mathbf{A}(\mathbf{s} - \boldsymbol{\alpha}), \mathbf{A}(\mathbf{s} - \boldsymbol{\alpha}) \rangle$$

$$= \sum_{i \in \mathcal{I}_B} \sum_{j \in \mathcal{I}_B} \sum_{\mathbf{y} \in \Sigma_i} \sum_{\bar{\mathbf{y}} \in \Sigma_j} (s(i,\mathbf{y}) - \alpha(i,\mathbf{y}))(s(j,\bar{\mathbf{y}}) - \alpha(j,\bar{\mathbf{y}})) \left\langle \frac{\boldsymbol{\Gamma}(\mathbf{t}_i, \mathbf{y}_i) - \boldsymbol{\Gamma}(\mathbf{t}_i, \mathbf{y})}{|E_i|}, \frac{\boldsymbol{\Gamma}(\mathbf{t}_j, \mathbf{y}_j) - \boldsymbol{\Gamma}(\mathbf{t}_j, \bar{\mathbf{y}})}{|E_j|} \right\rangle$$

$$= \sum_{i \in \mathcal{I}_B} \sum_{j \in \mathcal{I}_B} \sum_{\mathbf{y} \in \Sigma_i} \sum_{\bar{\mathbf{y}} \in \Sigma_j} (s(i,\mathbf{y}) - \alpha(i,\mathbf{y}))(s(j,\bar{\mathbf{y}}) - \alpha(j,\bar{\mathbf{y}})) \left\langle \frac{\boldsymbol{\Gamma}(\mathbf{t}_i, \mathbf{y}_i)}{|E_i|}, \frac{\boldsymbol{\Gamma}(\mathbf{t}_j, \mathbf{y}_j)}{|E_j|} \right\rangle \ldots$$

$$- \sum_{i \in \mathcal{I}_B} \sum_{j \in \mathcal{I}_B} \sum_{\mathbf{y} \in \Sigma_i} \sum_{\bar{\mathbf{y}} \in \Sigma_j} (s(i,\mathbf{y}) - \alpha(i,\mathbf{y}))(s(j,\bar{\mathbf{y}}) - \alpha(j,\bar{\mathbf{y}})) \left\langle \frac{\boldsymbol{\Gamma}(\mathbf{t}_i, \mathbf{y}_i)}{|E_i|}, \frac{\boldsymbol{\Gamma}(\mathbf{t}_j, \bar{\mathbf{y}})}{|E_j|} \right\rangle \ldots$$

$$- \sum_{i \in \mathcal{I}_B} \sum_{j \in \mathcal{I}_B} \sum_{\mathbf{y} \in \Sigma_i} \sum_{\bar{\mathbf{y}} \in \Sigma_j} (s(i,\mathbf{y}) - \alpha(i,\mathbf{y}))(s(j,\bar{\mathbf{y}}) - \alpha(j,\bar{\mathbf{y}})) \left\langle \frac{\boldsymbol{\Gamma}(\mathbf{t}_i, \mathbf{y})}{|E_i|}, \frac{\boldsymbol{\Gamma}(\mathbf{t}_j, \mathbf{y}_j)}{|E_j|} \right\rangle \ldots$$

$$+ \sum_{i \in \mathcal{I}_B} \sum_{j \in \mathcal{I}_B} \sum_{\mathbf{y} \in \Sigma_i} \sum_{\bar{\mathbf{y}} \in \Sigma_j} (s(i,\mathbf{y}) - \alpha(i,\mathbf{y}))(s(j,\bar{\mathbf{y}}) - \alpha(j,\bar{\mathbf{y}})) \left\langle \frac{\boldsymbol{\Gamma}(\mathbf{t}_i, \mathbf{y})}{|E_i|}, \frac{\boldsymbol{\Gamma}(\mathbf{t}_j, \bar{\mathbf{y}})}{|E_j|} \right\rangle \ldots$$

$$= \sum_{i \in \mathcal{I}_B} \sum_{j \in \mathcal{I}_B} \left\langle \frac{\boldsymbol{\Gamma}(\mathbf{t}_i, \mathbf{y}_i)}{|E_i|}, \frac{\boldsymbol{\Gamma}(\mathbf{t}_j, \mathbf{y}_j)}{|E_j|} \right\rangle \underbrace{\sum_{\mathbf{y} \in \Sigma_i} (s(i,\mathbf{y}) - \alpha(i,\mathbf{y}))}_{=0} \underbrace{\sum_{\bar{\mathbf{y}} \in \Sigma_j} (s(j,\bar{\mathbf{y}}) - \alpha(j,\bar{\mathbf{y}}))}_{=0} \ldots$$

$$- \sum_{i \in \mathcal{I}_B} \sum_{j \in \mathcal{I}_B} \sum_{\bar{\mathbf{y}} \in \Sigma_j} (s(j,\bar{\mathbf{y}}) - \alpha(j,\bar{\mathbf{y}})) \left\langle \frac{\boldsymbol{\Gamma}(\mathbf{t}_i, \mathbf{y}_i)}{|E_i|}, \frac{\boldsymbol{\Gamma}(\mathbf{t}_j, \bar{\mathbf{y}})}{|E_j|} \right\rangle \underbrace{\sum_{\mathbf{y} \in \Sigma_i} (s(i,\mathbf{y}) - \alpha(i,\mathbf{y}))}_{=0} \ldots$$

$$- \sum_{i \in \mathcal{I}_B} \sum_{j \in \mathcal{I}_B} \sum_{\mathbf{y} \in \Sigma_i} (s(i,\mathbf{y}) - \alpha(i,\mathbf{y})) \left\langle \frac{\boldsymbol{\Gamma}(\mathbf{t}_i, \mathbf{y})}{|E_i|}, \frac{\boldsymbol{\Gamma}(\mathbf{t}_j, \mathbf{y}_j)}{|E_j|} \right\rangle \underbrace{\sum_{\bar{\mathbf{y}} \in \Sigma_j} (s(j,\bar{\mathbf{y}}) - \alpha(j,\bar{\mathbf{y}}))}_{=0} \ldots$$

$$+ \sum_{i \in \mathcal{I}_B} \sum_{j \in \mathcal{I}_B} \sum_{\mathbf{y} \in \Sigma_i} \sum_{\bar{\mathbf{y}} \in \Sigma_j} (s(i,\mathbf{y}) - \alpha(i,\mathbf{y}))(s(j,\bar{\mathbf{y}}) - \alpha(j,\bar{\mathbf{y}})) \left\langle \frac{\boldsymbol{\Gamma}(\mathbf{t}_i, \mathbf{y})}{|E_i|}, \frac{\boldsymbol{\Gamma}(\mathbf{t}_j, \bar{\mathbf{y}})}{|E_j|} \right\rangle \ldots$$

$$= \sum_{i \in \mathcal{I}_B} \sum_{j \in \mathcal{I}_B} \sum_{\mathbf{y} \in \Sigma_i} \sum_{\bar{\mathbf{y}} \in \Sigma_j} (s(i,\mathbf{y}) - \alpha(i,\mathbf{y}))(s(j,\bar{\mathbf{y}}) - \alpha(j,\bar{\mathbf{y}})) \left\langle \frac{\boldsymbol{\Gamma}(\mathbf{t}_i, \mathbf{y})}{|E_i|}, \frac{\boldsymbol{\Gamma}(\mathbf{t}_j, \bar{\mathbf{y}})}{|E_j|} \right\rangle$$

$$= \sum_{i \in \mathcal{I}_B} \sum_{j \in \mathcal{I}_B} \sum_{\mathbf{y} \in \Sigma_i} \sum_{\bar{\mathbf{y}} \in \Sigma_j} (s(i,\mathbf{y}) - \alpha(i,\mathbf{y}))(s(j,\bar{\mathbf{y}}) - \alpha(j,\bar{\mathbf{y}})) \left\langle \frac{\boldsymbol{\Gamma}(\mathbf{t}_i, \mathbf{y})}{|E_i|}, \frac{\boldsymbol{\Gamma}(\mathbf{t}_j, \bar{\mathbf{y}})}{|E_j|} \right\rangle$$

$$
\begin{aligned}
&= \sum_{i\in\mathcal{I}_B}\sum_{j\in\mathcal{I}_B}\sum_{\mathbf{y}\in\Sigma_i}\sum_{\bar{\mathbf{y}}\in\Sigma_j}(s(i,\mathbf{y})-\alpha(i,\mathbf{y}))(s(j,\bar{\mathbf{y}})-\alpha(j,\bar{\mathbf{y}}))\sum_{(\sigma,\tau)\in E_i}\sum_{(\bar{\sigma},\bar{\tau})\in E_j}\frac{\langle\Gamma(\mathbf{t}_i^{\sigma\tau},\mathbf{y}^{\sigma\tau}),\Gamma(\mathbf{t}_j^{\bar{\sigma}\bar{\tau}},\bar{\mathbf{y}}^{\bar{\sigma}\bar{\tau}})\rangle}{|E_i|\cdot|E_j|}\\
&= \sum_{i\in\mathcal{I}_B}\sum_{j\in\mathcal{I}_B}\sum_{\mathbf{y}\in\Sigma_i}\sum_{\bar{\mathbf{y}}\in\Sigma_j}(s(i,\mathbf{y})-\alpha(i,\mathbf{y}))(s(j,\bar{\mathbf{y}})-\alpha(j,\bar{\mathbf{y}}))\times\dots\\
&\qquad\sum_{(\sigma,\tau)\in E_i}\sum_{(\bar{\sigma},\bar{\tau})\in E_j}\frac{\operatorname{sign}(t_{i\sigma}-t_{i\tau})\operatorname{sign}(t_{j\bar{\sigma}}-t_{j\bar{\tau}})(\lambda(y_\sigma,\bar{y}_\sigma)-\lambda(y_\sigma,\bar{y}_\tau)-\lambda(y_\tau,\bar{y}_\sigma)+\lambda(y_\tau,\bar{y}_\tau))}{|E_i|\cdot|E_j|}\\
&= \sum_{i\in\mathcal{I}_B}\sum_{j\in\mathcal{I}_B}\sum_{\mathbf{y}\in\Sigma_i}\sum_{\bar{\mathbf{y}}\in\Sigma_j}(s(i,\mathbf{y})-\alpha(i,\mathbf{y}))(s(j,\bar{\mathbf{y}})-\alpha(j,\bar{\mathbf{y}}))\left(\frac{\operatorname{sign}(\Delta\mathbf{t}_i)}{|E_i|}\right)^T\mathbf{P}_i\mathbf{\Lambda}_{\mathbf{y}\bar{\mathbf{y}}}\mathbf{P}_j^T\left(\frac{\operatorname{sign}(\Delta\mathbf{t}_j)}{|E_j|}\right),
\end{aligned}
$$

with $\mathbf{P}_i \in \{-1,0,1\}^{|E_i|\times L}$ being a matrix encoding the edges such that:

$$
[\mathbf{P}_i]_{((\sigma,\tau),\cdot)} = \left[0,\dots,0,\underbrace{1}_{\sigma},0,\dots,0,\underbrace{-1}_{\tau},0,\dots,0\right] \qquad \forall(\sigma,\tau)\in E_i,
$$

and $\lambda:\mathcal{Y}\times\mathcal{Y}\rightarrow\mathbb{R}$ being a kernel function to compute the similarly between to molecules (see Methods "Molecule feature representations"), and $\mathbf{\Lambda}_{\mathbf{y}\bar{\mathbf{y}}}\in\mathbb{R}^{|V_i|\times|V_j|}$ being a kernel matrix between the molecules associated with the label sequences $\mathbf{y}\in\Sigma_i$ and $\bar{\mathbf{y}}\in\Sigma_j$, $\operatorname{sign}(\Delta\mathbf{t}_i) = (\operatorname{sign}(t_{i\sigma}-t_{i\tau}))_{(\sigma,\tau)\in E_i}\in\{-1,0,1\}^{|E_i|}$ being a vector collecting the signs of the retention time differences.

#### 1.3 Scoring function $F$

$$
\begin{aligned}
F(\mathbf{y}\,|\,\mathbf{x}_i,\mathbf{t}_i,\mathbf{A}\boldsymbol{\alpha},G_i) &= \frac{\boldsymbol{\theta}(\mathbf{x}_i,\mathbf{y})}{|V_i|}+\frac{\langle\mathbf{\Gamma}(\mathbf{t}_i,\mathbf{y}),\mathbf{A}\boldsymbol{\alpha}\rangle}{|E_i|}\\
&= \frac{1}{|V_i|}\sum_{\sigma\in V_i}\theta(x_{i\sigma},y_\sigma)+\frac{1}{|E_i|}\sum_{(\sigma,\tau)\in E_i}\langle\Gamma(\mathbf{t}_i^{\sigma\tau},\mathbf{y}^{\sigma\tau}),\mathbf{A}\boldsymbol{\alpha}\rangle\\
&= \frac{1}{|V_i|}\sum_{\sigma\in V_i}\theta(x_{i\sigma},y_\sigma)+\frac{1}{|E_i|}\sum_{(\sigma,\tau)\in E_i}\operatorname{sign}(t_{i\sigma}-t_{i\tau})\langle\phi(y_{i\sigma})-\phi(y_{i\tau}),\mathbf{y}^{\sigma\tau}),\mathbf{A}\boldsymbol{\alpha}\rangle\\
&= \frac{1}{|V_i|}\sum_{\sigma\in V_i}\theta(x_{i\sigma},y_\sigma)+\dots\\
&\qquad\frac{1}{|E_i|}\left(\sum_{(\sigma,\tau)\in E_i}\operatorname{sign}(t_{i\sigma}-t_{i\tau})\underbrace{\langle\phi(y_{i\sigma}),\mathbf{A}\boldsymbol{\alpha}\rangle}_{\equiv h(y_{i\sigma}\,|\,\mathbf{A}\boldsymbol{\alpha})}-\sum_{(\sigma,\tau)\in E_i}\operatorname{sign}(t_{i\sigma}-t_{i\tau})\langle\phi(y_{i\tau}),\mathbf{A}\boldsymbol{\alpha}\rangle\right)\\
&= \frac{1}{|V_i|}\sum_{\sigma\in V_i}\theta(x_{i\sigma},y_\sigma)+\sum_{(\sigma,\tau)\in E_i}\dots\\
&\qquad\sum_{j=1}^{N}\sum_{\bar{\mathbf{y}}\in\Sigma_j}\alpha(j,\bar{\mathbf{y}})\sum_{(\bar{\sigma},\bar{\tau})\in E_j}\left\langle\frac{\Gamma(\mathbf{t}_i^{\sigma\tau},\mathbf{y}^{\sigma\tau})}{|E_i|},\frac{\Gamma(\mathbf{t}_j^{\bar{\sigma}\bar{\tau}},\bar{\mathbf{y}}_j^{\bar{\sigma}\bar{\tau}})-\Gamma(\mathbf{t}_j^{\bar{\sigma}\bar{\tau}},\bar{\mathbf{y}}^{\bar{\sigma}\bar{\tau}})}{|E_j|}\right\rangle\\
&= \frac{1}{|V_i|}\sum_{\sigma\in V_i}\theta(x_{i\sigma},y_\sigma)+\sum_{(\sigma,\tau)\in E_i}\Bigg(\dots\\
&\qquad\sum_{j=1}^{N}\sum_{\bar{\mathbf{y}}\in\Sigma_j}\alpha(j,\bar{\mathbf{y}})\sum_{(\bar{\sigma},\bar{\tau})\in E_j}\left\langle\frac{\Gamma(\mathbf{t}_i^{\sigma\tau},\mathbf{y}^{\sigma\tau})}{|E_i|},\frac{\Gamma(\mathbf{t}_j^{\bar{\sigma}\bar{\tau}},\bar{\mathbf{y}}_j^{\bar{\sigma}\bar{\tau}})}{|E_j|}\right\rangle-\dots
\end{aligned}
$$

$$
\begin{aligned}
&\sum_{j=1}^{N}\sum_{\bar{\mathbf{y}}\in\Sigma_j}\alpha(j,\bar{\mathbf{y}})\sum_{(\bar\sigma,\bar\tau)\in E_j}\left\langle \frac{\Gamma(\mathbf{t}_i^{\sigma\tau},\mathbf{y}^{\sigma\tau})}{|E_i|}, \frac{\Gamma(\mathbf{t}_j^{\bar\sigma\bar\tau},\bar{\mathbf{y}}^{\bar\sigma\bar\tau})}{|E_j|}\right\rangle\Bigg)\\
= &\frac{1}{|V_i|}\sum_{\sigma\in V_i}\theta(x_{i\sigma},y_\sigma) + \sum_{(\sigma,\tau)\in E_i}\dots\\
&\sum_{j=1}^{N}\sum_{\bar{\mathbf{y}}\in\Sigma_j}\alpha(j,\bar{\mathbf{y}})\sum_{(\bar\sigma,\bar\tau)\in E_j}\left\langle \frac{\operatorname{sign}(t_{i\sigma}-t_{i\tau})(\phi(y_{i\sigma})-\phi(y_{i\tau}))}{|E_i|}, \frac{\Gamma(\mathbf{t}_j^{\bar\sigma\bar\tau},\bar{\mathbf{y}}_j^{\bar\sigma\bar\tau})-\Gamma(\mathbf{t}_j^{\bar\sigma\bar\tau},\bar{\mathbf{y}}^{\bar\sigma\bar\tau})}{|E_j|}\right\rangle\\
= &\frac{1}{|V_i|}\sum_{\sigma\in V_i}\theta(x_{i\sigma},y_\sigma) + \sum_{(\sigma,\tau)\in E_i}\frac{\operatorname{sign}(t_{i\sigma}-t_{i\tau})}{|E_i|}\times\dots\\
&\sum_{j=1}^{N}\sum_{\bar{\mathbf{y}}\in\Sigma_j}\alpha(j,\bar{\mathbf{y}})\sum_{(\bar\sigma,\bar\tau)\in E_j}\left\langle \phi(y_{i\sigma})-\phi(y_{i\tau}), \frac{\Gamma(\mathbf{t}_j^{\bar\sigma\bar\tau},\bar{\mathbf{y}}_j^{\bar\sigma\bar\tau})-\Gamma(\mathbf{t}_j^{\bar\sigma\bar\tau},\bar{\mathbf{y}}^{\bar\sigma\bar\tau})}{|E_j|}\right\rangle\\
= &\frac{1}{|V_i|}\sum_{\sigma\in V_i}\theta(x_{i\sigma},y_\sigma) + \sum_{(\sigma,\tau)\in E_i}\frac{\operatorname{sign}(t_{i\sigma}-t_{i\tau})}{|E_i|}\times\Bigg(\dots\\
&\sum_{j=1}^{N}\sum_{\bar{\mathbf{y}}\in\Sigma_j}\alpha(j,\bar{\mathbf{y}})\sum_{(\bar\sigma,\bar\tau)\in E_j}\left\langle \phi(y_{i\sigma}), \frac{\Gamma(\mathbf{t}_j^{\bar\sigma\bar\tau},\bar{\mathbf{y}}_j^{\bar\sigma\bar\tau})-\Gamma(\mathbf{t}_j^{\bar\sigma\bar\tau},\bar{\mathbf{y}}^{\bar\sigma\bar\tau})}{|E_j|}\right\rangle - \dots\\
&\sum_{j=1}^{N}\sum_{\bar{\mathbf{y}}\in\Sigma_j}\alpha(j,\bar{\mathbf{y}})\sum_{(\bar\sigma,\bar\tau)\in E_j}\left\langle \phi(y_{i\tau}), \frac{\Gamma(\mathbf{t}_j^{\bar\sigma\bar\tau},\bar{\mathbf{y}}_j^{\bar\sigma\bar\tau})-\Gamma(\mathbf{t}_j^{\bar\sigma\bar\tau},\bar{\mathbf{y}}^{\bar\sigma\bar\tau})}{|E_j|}\right\rangle\Bigg)
\end{aligned}
$$

Supplementary Table 1: MassBank (MB) information used to group the records. Two MassBank records are considered to belong to the same MB-subset in our experiments, if all properties listed in the table are equal between them. See `https://github.com/MassBank/MassBank-web/blob/main/Documentation/MassBankRecordFormat.md` for a more comprehensive description of the MassBank records' fields.

| **Property** | **Description** | **Example** |
|---|---|---|
| `contributor` | Contributor who uploaded a MassBank record | BGC_Munich |
| `accession prefix` | 2-3 character long prefix further specifying the records of a contributor | EA, EQ |
| `instrument_type` | General type of instrument used for the LC-MS analysis | LC-ESI-QTOF |
| `ion_mode` | MS Ionization mode | negative |
| `instrument` | Commercial name and manufacturer of the MS instrument | Bruker maXis Impact |
| `fragmentation_mode` | Fragmentation method used for dissociation or fragmentation | CID |
| `column_name` | Commercial name and manufacturer of the LC instrument | Symmetry C18 Column, Waters |
| `column_temperature` | Static column temperature in LC-MS | 40 C |
| `flow_gradient` | Gradient of mobile phases in LC-MS | 0min:5%, 24min:95% (acetonitrile) |
| `flow_rate` | Flow Rate of liquid phase in LC | 300 uL/min |
| `solvent_A` | Chemical composition of buffer solution (A) | H2O(0.1%HCOOH) |
| `solvent_B` | Chemical composition of buffer solution (B) | CH3CN(0.1%HCOOH) |

Supplementary Table 2: Training and evaluation dataset sizes in our experiments. We provide the number (#) of ($MS^2$, RT)-tuples used for the generation of training and evaluation LC-$MS^2$ experiments. For the ALLDATA setup the training and evaluation tuple-set is equal. The number of evaluation LC-$MS^2$ experiments depends on the number of available evaluation tuples.

| | **ALLDATA** | | **ONLYSTEREO** | | |
|---|---|---|---|---|---|
| **MB-subset** | **#Tuples** | **#Exp.** | **#Tuples (train.)** | **#Tuples (eval.)** | **#Exp.** |
| AC_003 | 179 | 15 | 172 | 157 | 15 |
| AU_000 | 168 | 15 | 146 | 23 | - |
| AU_002 | 746 | 14 | 578 | 172 | 15 |
| AU_003 | 90 | 15 | 77 | 21 | - |
| BML_000 | 170 | 15 | 77 | 24 | - |
| BML_001 | 250 | 15 | 125 | 33 | 1 |
| BS_000 | 216 | 15 | 205 | 135 | 15 |
| CE_001 | 39 | 1 | 30 | 19 | - |
| EA_000 | 141 | 15 | 118 | 19 | - |
| EA_001 | 147 | 15 | 126 | 19 | - |
| EA_002 | 301 | 6 | 240 | 56 | 1 |
| EA_003 | 307 | 6 | 246 | 57 | 1 |
| EQ_001 | 86 | 15 | 68 | 28 | - |
| EQ_003 | 92 | 15 | 64 | 6 | - |
| EQ_004 | 181 | 15 | 127 | 51 | 1 |
| EQ_006 | 211 | 15 | 138 | 15 | - |
| ET_002 | 50 | 1 | 29 | 2 | - |
| KW_000 | 55 | 1 | 43 | 4 | - |
| LQB_000 | 301 | 6 | 271 | 270 | 5 |
| LU_000 | 358 | 7 | 311 | 50 | 1 |
| LU_001 | 567 | 11 | 472 | 101 | 15 |
| NA_003 | 97 | 15 | 91 | 73 | 1 |
| PR_000 | 709 | 14 | 131 | 21 | - |
| PR_002 | 911 | 18 | 391 | 250 | 15 |
| RP_000 | 69 | 1 | 55 | 35 | 1 |
| RP_001 | 150 | 15 | 119 | 73 | 1 |
| SM_000 | 161 | 15 | 136 | 12 | - |
| SM_001 | 357 | 7 | 280 | 30 | 1 |
| UF_002 | 149 | 15 | 124 | 18 | - |
| UF_003 | 140 | 15 | 115 | 15 | - |
| UT_000 | 318 | 6 | 294 | 293 | 5 |
| Total | 7716 | 354 | 5399 | 2082 | 94 |

Supplementary Table 3: Meta-information for each MassBank (MB) subset used in our study. This encompoasses the LC- and $MS^2$-conditions of each MB-subset.

THIS TABLE IS PROVIDED IN A SEPARATE FILE: `massbank_subsets_meta_data.csv` AND ALSO AVAILABLE ON ZENODO: [7]

Supplementary Table 4: Candidate score aggregation when collapsing by InChIKey-1 or InChIKey. Illustration of the score aggregation when collapsing the candidates by either InChIKey-1 (first block, ALLDATA) or InChIKey (ONLYSTEREO). For each candidate group, defined by the InChIKey part, the maximum score computed and a reduced candidate set is returned. In our experiments the score can be Only-MS$^2$, LC-MS$^2$Struct, etc.

| CID | InChIKey | Score |
|---|---|---|
| 29921449 | KZWDYKRBPAXQDR-KVQBGUIXSA-N | 0.98 |
| 97988234 | KZWDYKRBPAXQDR-SRQIZXRXSA-N | 0.95 |
| 70367116 | RESQOVGTIWHTLH-IDOCCRKTSA-N | 0.78 |
| 153640319 | RESQOVGTIWHTLH-IDOCCRKTSA-N | 0.79 |
| 101065727 | IHEGXRGRQBFUTL-KVQBGUIXSA-N | 0.87 |
| 88274880 | JPRLCMCVIAOOIT-DQSPEZDDSA-N | 0.92 |
| 196710 | JPRLCMCVIAOOIT-OBXARNEKSA-N | 0.96 |
| 14806063 | JPRLCMCVIAOOIT-UHFFFAOYSA-N | 0.97 |

Maximum score per InChIKey-1 group ⟹

| InChIKey-1 | Score |
|---|---|
| KZWDYKRBPAXQDR | 0.98 |
| RESQOVGTIWHTLH | 0.79 |
| IHEGXRGRQBFUTL | 0.87 |
| JPRLCMCVIAOOIT | 0.97 |

Maximum score per InChIKey group ⇓

| InChIKey | Score |
|---|---|
| KZWDYKRBPAXQDR-KVQBGUIXSA-N | 0.98 |
| KZWDYKRBPAXQDR-SRQIZXRXSA-N | 0.95 |
| RESQOVGTIWHTLH-IDOCCRKTSA-N | 0.79 |
| IHEGXRGRQBFUTL-KVQBGUIXSA-N | 0.87 |
| JPRLCMCVIAOOIT-DQSPEZDDSA-N | 0.92 |
| JPRLCMCVIAOOIT-OBXARNEKSA-N | 0.96 |
| JPRLCMCVIAOOIT-UHFFFAOYSA-N | 0.97 |

Supplementary Table 5: Median candidate set size for the MassBank (MB) subsets. The table shows the median number of molecular candidates per MB-subset used in our experiments. In the ALLDATA setup the candidates are identified by their InChIKey first block, where as for the Only-MS$^2$ setup the full InChIKey is used. The candidate number is computed based on the MB records which are used in the simulated LC-MS$^2$ experiments. For ONLYSTEREO, some MB-subsets are not used in the evaluation, and therefore their candidate set size is omitted (-).

| **MB-subset** | **ALLDATA** | **ONLYSTEREO** |
|---|---:|---:|
| AC_003 | 305 | 384 |
| AU_000 | 269 | - |
| AU_002 | 1018.5 | 1434.5 |
| AU_003 | 1297 | - |
| BML_000 | 689 | - |
| BML_001 | 1013.5 | 1688 |
| BS_000 | 429 | 258 |
| CE_001 | 819 | - |
| EA_000 | 771 | - |
| EA_001 | 570 | - |
| EA_002 | 1373 | 1239 |
| EA_003 | 1306 | 1097 |
| EQ_001 | 425 | - |
| EQ_003 | 759.5 | - |
| EQ_004 | 872 | 1027 |
| EQ_006 | 1045 | - |
| ET_002 | 4957 | - |
| KW_000 | 2010 | - |
| LQB_000 | 73 | 106 |
| LU_000 | 533 | 362.5 |
| LU_001 | 998 | 751 |
| NA_003 | 1024 | 1608 |
| PR_000 | 109 | - |
| PR_002 | 228 | 636 |
| RP_000 | 760 | 1015 |
| RP_001 | 658 | 723 |
| SM_000 | 312 | - |
| SM_001 | 800 | 1095.5 |
| UF_002 | 1498 | - |
| UF_003 | 1392.5 | - |
| UT_000 | 56 | 93 |
